# Peptipedia v2.0: A peptide sequence database and user-friendly web platform. A major update

**DOI:** 10.1101/2024.07.11.603053

**Authors:** Gabriel Cabas-Mora, Anamaría Daza, Nicole Soto-García, Valentina Garrido, Diego Alvarez, Marcelo Navarrete, Lindybeth Sarmiento-Varón, Julieta H. Sepúlveda Yañez, Mehdi D. Davari, Frederic Cadet, Álvaro Olivera-Nappa, Roberto Uribe-Paredes, David Medina-Ortiz

**Affiliations:** Departamento de Ingeniería En Computación, Universidad de Magallanes, Avenida Bulnes 01855, Punta Arenas, Chile; Centre for Biotechnology and Bioengineering, CeBiB, Universidad de Chile, Beauchef 851, Santiago, Chile; Centro Asistencial de Docencia e Investigación, CADI, Universidad de Magallanes. Av. Los Flamencos 01364, Punta Arenas, Chile; Escuela de Medicina, Universidad de Magallanes, Avenida Bulnes 01855, Punta Arenas, Chile; Facultad de Ciencias de la Salud, Universidad de Magallanes, Avenida Bulnes 01855, Punta Arenas, Chile; Department of Bioorganic Chemistry, Leibniz Institute of Plant Biochemistry, Weinberg 3, 06120, Halle, Germany; PEACCEL, Artificial Intelligence Department, AI for Biologics, 75013, Paris, France

**Keywords:** Peptide databases, Enrichment analysis, binary classification models, protein language models, bioinformatics tools, retention time predictive models

## Abstract

In recent years, peptides have gained significant relevance due to their therapeutic properties. The surge in peptide production and synthesis has generated vast amounts of data, enabling the creation of comprehensive databases and information repositories. Advances in sequencing techniques and artificial intelligence have further accelerated the design of tailor-made peptides. However, leveraging these techniques requires versatile and continuously updated storage systems, along with tools that facilitate peptide research and the implementation of machine learning for predictive systems. This work introduces Peptipedia v2.0, one of the most comprehensive public repositories of peptides, supporting biotechnological research by simplifying peptide study and annotation. Peptipedia v2.0 has expanded its collection by over 45% with peptide sequences that have reported biological activities. The functional biological activity tree has been revised and enhanced, incorporating new categories such as cosmetic and dermatological activities, molecular binding, and anti-ageing properties. Utilizing protein language models and machine learning, more than 90 binary classification models have been trained, validated, and incorporated into Peptipedia v2.0. These models exhibit average sensitivities and specificities of 0.877 ± 0.0530 and 0.873 ±0.054, respectively, facilitating the annotation of more than 3.6 million peptide sequences with unknown biological activities, also registered in Peptipedia v2.0. Additionally, Peptipedia v2.0 introduces description tools based on structural and ontological properties and user-friendly machinelearning tools to facilitate the application of machine-learning strategies to study peptide sequences. Peptipedia v2.0 is accessible under the Creative Commons CC BY-NC-ND 4.0 license at https://peptipedia.cl/.

## 1 Introduction

Peptides are versatile biomolecules that can be synthetic or found in natural sources and are attractive candidates for therapeutic applications (Lau and Dunn, 2018; Lien and Lowman, 2003; Wang et al., 2022). Peptides play crucial roles in numerous biological processes. Their functions are diverse, as structural components, enzymatic inhibitors, cell-penetrating agents, hormones, host defence molecules, and neurotransmitters (Wang et al., 2022). Additionally, peptides act as cell surface receptors (Taylor, 2000) and are integral to drug delivery applications (Khan et al., 2021).

Peptide drugs present several potential advantages over traditional small-molecule drugs, such as increased selectivity, affinity, efficacy, and safety, along with reduced toxicity and immunogenicity (Apostolopoulos et al., 2021). However, their widespread clinical application is hindered by challenges, including a short half-life, limited oral bioavailability, and susceptibility to plasma degradation (Goles et al., 2024).

The therapeutic market started 100 years ago using insulin to treat type 1 diabetes Sims et al., 2021; Goeddel et al., 1979. To date, over 100 peptide drugs are currently available in the global market for various diseases, including HIV infection (human immunodeficiency virus), chronic pain, metabolic disorders, and infectious diseases (Henninot et al., 2018; Wang et al., 2022; Lee et al., 2019). The global therapeutic peptides market reached a value of US$42.05 billion in 2022 (GVR Report Cover, 2023). Due to research efforts, developed technology, and the investment of pharmaceutical companies, it is expected that peptide drug discovery and development will continue to expand in the upcoming years, foreseen to reach US$68.6 billion by 2030 (with a compound annual growth rate (CAGR) 6.3% from 2023 to 2030) (Muttenthaler et al., 2021; Lau and Dunn, 2018; Wan et al., 2022; GVR Report Cover, 2023).

The scarcity of structured and specialised peptide repositories motivated the implementation of the first version of the Peptipedia database and its user-friendly web platform (Quiroz et al., 2021). Peptipedia v1.0 collect over 92,000 peptide sequences and more than 75,000 peptides with reported biological activity from 30 data sources. The Peptipedia v1.0 web platform also included a functional physicochemical properties estimator for peptide sequences, statistical amino acid sequence characterization, and bioinformatics tools like sequence alignment to facilitate the study of peptide sequences. Furthermore, Peptipedia v1.0 implemented machine learning-based binary classification models to identify potential biological activities like antimicrobial, antiviral, and antibacterial.

Before Peptipedia v1.0, peptide information was widely spread in diverse databases such as DBAASP, Erop-Moscow, LAMP, DRAMP, SATPdb (Pirtskhalava et al., 2020a; Zamyatnin, 1991; Zhao et al., 2013; Kang et al., 2019; Singh et al., 2016). Peptide-related databases comprehend from a very specialised topic, i.e., blood-brain barrier peptides (Van Dorpe et al., 2012), quorum sensing peptides (Wynendaele et al., 2013), anti-angiogenic peptides predictor (Ramaprasad et al., 2015), and bacteriocin peptides (Hammami et al., 2010) to general databases, i.e., the Universal Protein Resource (UniProt) (uni, 2023), LAMP2, a generic antimicrobial peptide database (Zhao et al., 2013), DRAMP, a data repository of antimicrobial peptides (Kang et al., 2019), and SATPdb, a database of structurally annotated therapeutic peptides (Singh et al., 2016).

Despite the availability of numerous databases, integrating information between them remains challenging. Peptides are usually classified by just one functional biological activity. However, moonlight peptides have two or more known activities within the same domain (Jeffery, 1999), and identifying the potential multiple activities of a peptide is relevant for biotechnological and pharmaceutical industries (Singh and Bhalla, 2020; Zanzoni et al., 2019).

The Peptipedia database is a knowledge-enhancing peptide research tool that extracts, consolidates, organises, and curates information from multiple databases. While Peptipedia v1.0 is a powerful resource for peptide research, it requires addressing several issues. Firstly, the number of records could be significantly increased, and the information related to the peptides could be expanded. Secondly, the current classification of biological activities is limited, often categorising many peptide activities as *“other”*. Finally, the absence of an automatic updating system for newly generated data hinders its full potential. By addressing these issues, Peptipedia can be enhanced, providing researchers with a more comprehensive and dynamic knowledge platform with data integration and harmonization across platforms.

This work presents a significant update to the Peptipedia database, incorporating over 3.8 million peptide sequences from more than 70 data sources. Over 100,000 peptides are documented as having functional biological activity based on literature. The categories and subcategories of the biological activity tree were meticulously examined and reorganized, with new activities such as anti-ageing, cytokine, and molecular binding included. An automatic update system was also implemented, and the advanced search and downloading functionalities were enhanced.

In Peptipedia v2.0, various types of information have been linked to each peptide sequence, including functional domains (Mistry et al., 2021), gene ontology annotations (Ashburner et al., 2000; Aleksander et al., 2023), secondary structure information (Wang et al., 2016), and 3D structures available through PDB (Berman et al., 2000a) and AlphaFold (Jumper et al., 2021; Van Dorpe et al., 2012) databases.

Moreover, Peptipedia v2.0 incorporates data on physicochemical property descriptors, patents, and previous publications. Bioinformatic enrichment analysis and statistical evaluation tools have also been integrated. Furthermore, the functional classification models have been updated, enhancing the precision of individual models and improving their generalization. These trained classification models evaluated over 3.7 million peptide sequences, predicting their functional biological activities.

Finally, Peptipedia v2.0 includes machine learning tools for users to analyze their datasets and build their models. These tools encompass protein language models and classical amino acid coding methods for numerical representation strategies, predictive model training through traditional supervised learning algorithms, pattern recognition using unsupervised learning approaches, and sequence similarity networks combined with community detection techniques for pattern recognition. With these enhancements, users can gain deeper insights and uncover novel characteristics from their datasets. Consequently, all these new tools and updates establish Peptipedia v2.0 as one of the most comprehensive, thoroughly processed, and tractable repositories of peptides currently available.

## 2 Methodologies and implementation strategies

### 2.1 Collecting and processing peptide sequences

This work searched databases, datasets, and public repositories related to the study of peptides to update and increase the number of records in the Peptipedia v2.0 database. Different keywords were used to collect the data sources in Google Scholar, like *peptides, AMPs, neuropeptides, anticancer peptides, nutraceutical peptides*, and *signal peptides*. Besides, generic protein databases such as UniProtKB (uni, 2023) and RCSB PDB (Berman et al., 2000b) were incorporated (See section S1 of Supplementary Information for more details).

Information related to the characteristics of the reported peptide sequences, including their biological activities, descriptions, experimental information, and related publications or patents, was downloaded from all data sources.

Then, Python scripts were implemented to process the raw data downloaded from the data sources and transform the information for loading into the Peptipedia v2.0.

A length filter was made, containing only peptides with a length equal to or less than 150 residues and higher than three residues. Also, the collected peptides were classified as canonical (with only the 20 natural amino acids) or non-canonical peptides. Then, a semantic analysis was generated to recognise the biological activity of the peptide sequence through the available description in the data sources.

Finally, a loader Python script was implemented to load the register in the Peptipedia v2.0, developing a scalable ETL strategy (Extract, Transformation, and Load) for each utilised data source.

### 2.2 Peptide descriptions and enrichment analysis implementation

Each peptide incorporated in the Peptipedia v2.0 was characterised using the peptide descriptor service and enrichment analysis system.

First, the modlAMP tool (Müller et al., 2017) was used to calculate the physicochemical properties of canonical peptide sequences. Then, the standalone version of RaptorX-Property tool (Wang et al., 2016) was employed to predict the secondary structure of all records available in the Peptipedia v2.0.

Through homology mechanisms, enrichment analysis was performed for all canonical peptides using the MetaStudent tool (Hamp et al., 2013). MetaStudent allows the assignment of gene ontology terms (GO) from different sources, such as molecular function, cellular localization, and biological process.

Once the gene ontology terms prediction was completed, the results were filtered using the data returned probability score, selecting only results with a probability higher than 0.5 as elements to incorporate as a descriptor of the analysed peptide sequence. Finally, PfamScan tool (Mistry et al., 2020) was used to estimate protein domain families for all peptides registered in the Peptipedia v2.0.

### 2.3 Training predictive models

This work implements binary classification models to identify peptide sequences’ functional biological activities by combining embedding representation through pre-trained models with classic supervised learning algorithms (Medina-Ortiz Sr et al., 2024).

A binary dataset was constructed for each biological activity identified in the Peptipedia v2.0. The following steps were used to build the datasets: i) Peptides exhibiting the target activity were collected to generate positive examples, ii) Peptides without the target activity were collected to create the negative examples, iii) An undersampling strategy was applied to balance the dataset by randomly removing negative examples, and iv) Different pre-trained models were then used to generate embeddings, representing peptide sequences numerically for training the classification models (Dallago et al., 2021) (See more details in Section S2 of Supplementary Materials).

Activities with fewer than 50 peptides were excluded because they are classified as *Low-N* datasets, and more specialized strategies, such as transfer learning, semi-supervised approaches, and contrastive learning methods, are required to train predictive models using these types of datasets (Biswas et al., 2021).

Each dataset built is divided into training and validation, using an 80:20 proportion. Nine supervised learning algorithms, including Decision Tree, Random Forest, and XGBoost, were used to train classification models using the training dataset. A *K*-fold cross-validation (*k* = 10) was performed to prevent overfitting. The validation dataset was then used to assess model performance using classical metrics such as precision, recall, accuracy, and F-score (Medina-Ortiz et al., 2020a).

The combination of supervised learning algorithm and embedding representation is selected using the performances obtained during the training and validation process, applying the following criteria: i) highest performance during the training stage, ii) highest performance during the validation stage, and iii) lowest differences between training and validation to reduce the overfitting.

Finally, the models were exported in joblib format for integration into the Peptipedia v2.0.

### 2.4 Implementation strategies and availability system

The database available in Peptipedia v2.0 was designed based on a relational schema. PostgreSQL manages all operations over the database. All queries performed to the database are managed through an application programming interface (API) implemented using the Flask framework version 2.3.1 and SQLAlchemy version 2.0.29.

The ETL system, description process, and enrichment analysis strategies were implemented using Python version 3.9.17. Moreover, each tool and service available in Peptipedia v2.0 was implemented using Python programming version 3.9.17.

All binary classification models were implemented using Python v3.9.17 and the supervised learning module available on the DMAKit Python library (Medina-Ortiz et al., 2020b). Moreover, the embedding representation for the peptide sequences was performed using the bio-embedding library (Dallago et al., 2021).

Finally, the user-friendly web platform in Peptipedia v2.0 was implemented using the React framework as the front end. The system was deployed using Podman into a public server with AlmaLinux 9 as the operative system. Its hardware characteristics are eight vCPU Cores, 64 GB RAM, 500 GB NVMe, and 32 TB Traffic.

## 3 Results and discussions

Peptipedia v2 is a user-friendly web platform designed to facilitate the study of peptide sequences using advanced bioinformatics tools and machine-learning strategies. This updated version boasts a database of over 100,000 peptides sourced from more than 70 databases. Various physicochemical and thermodynamic properties characterize each peptide.

New features in Peptipedia v2 include enrichment analysis through Gene Ontology terms, secondary structure predictions, and functional domain evaluations, providing more comprehensive information on each peptide sequence. The functional biological activity tree has been enhanced to include more specific cosmetics, dermatology, taste, and molecular binding activities.

Additionally, the platform now implements over 90 binary classification models for biological activity classification, utilizing embedding representations from pre-trained models and traditional supervised learning algorithms. Numerous tools, such as advanced search capabilities, peptide sequence downloading, physicochemical characterization, and functional biological activity classification, have been updated or newly incorporated.

Moreover, Peptipedia v2 includes an integrative machine-learning pipeline to streamline the development of sequence-based predictive models and pattern recognition. An overview of Peptipedia v2.0 is presented in Figure 1, highlighting the most relevant data sources, peptide sequence information, bioinformatics tools, and machine learning applications.

**Figure 1:**
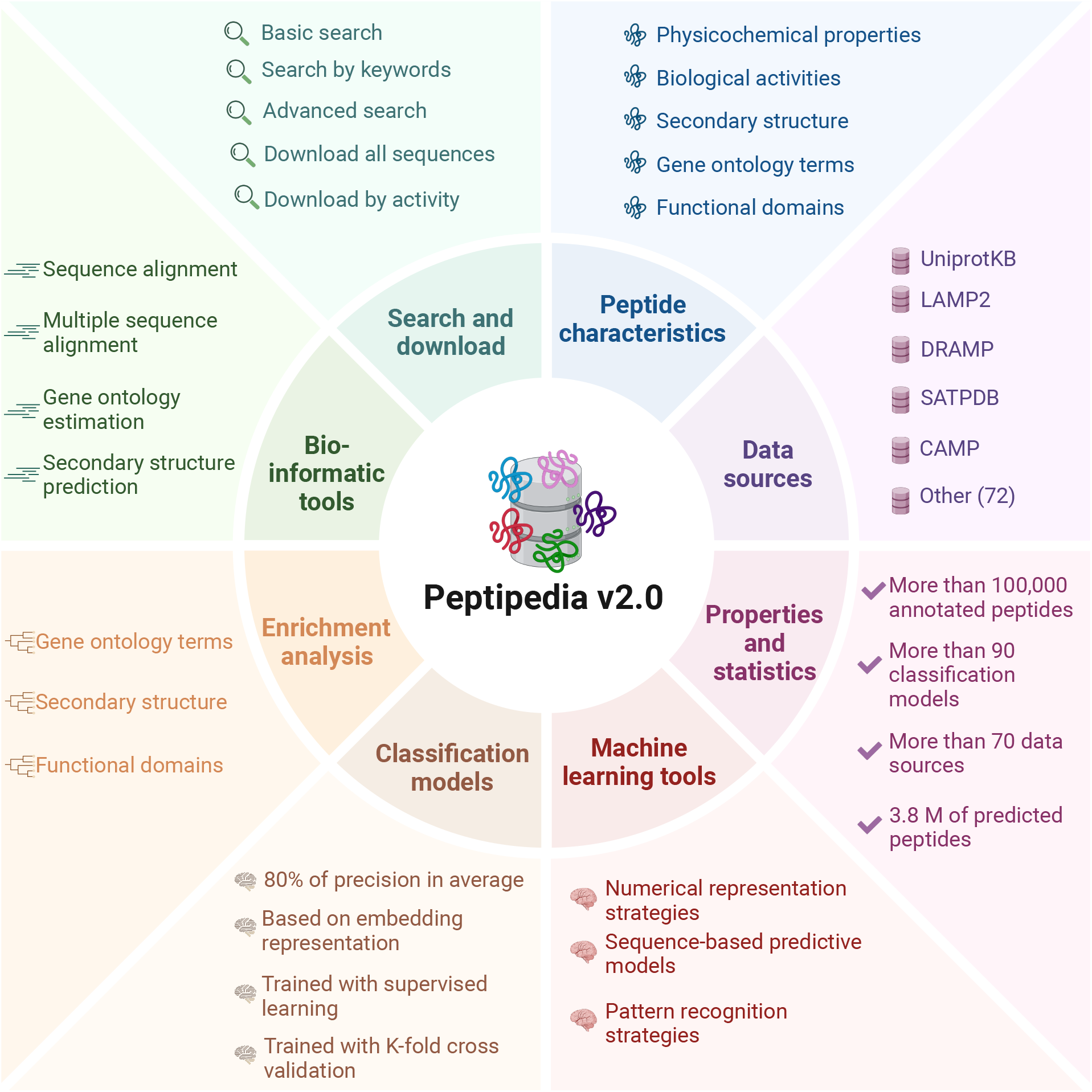
Overview of Peptipedia v2.0. This new version of Peptipedia includes more than 100,000 peptides registered with functional biological activities, extracted from more than 70 data sources. Different physicochemical properties and enrichment analyses were addressed to characterize the peptides collected. The services and functionalities were updated, incorporating gene ontology and functional domain predictions, secondary structure evaluation, and physicochemical properties estimation. Besides, more than 90 functional biological activity classification models were implemented, combining embedding representation through pre-trained models and supervised learning algorithms. Finally, customizable machine learning pipelines could be implemented by employing the machine learning tools in Peptipedia v2.0 to facilitate the application of ML techniques to study peptide sequences.

### 3.1 New data sources, biological activity tree, and peptide sequences

Peptipedia v2.0 includes 76 data sources, comprising databases, repositories, and datasets (see Section S1 of Supplementary Materials for more details). These data sources are dedicated to collecting and retrieving peptide sequences alongside their corresponding biological activities. The data sources used in this Peptipedia v2.0 database increased by over 100% in the number of data sources compared to the initial version of Peptipedia (Quiroz et al., 2021). A comprehensive update to the biological activity tree has been executed alongside expanding data sources. Figure 2 summarizes the updated functional biological activity tree in Peptipedia v2.0.

**Figure 2:**
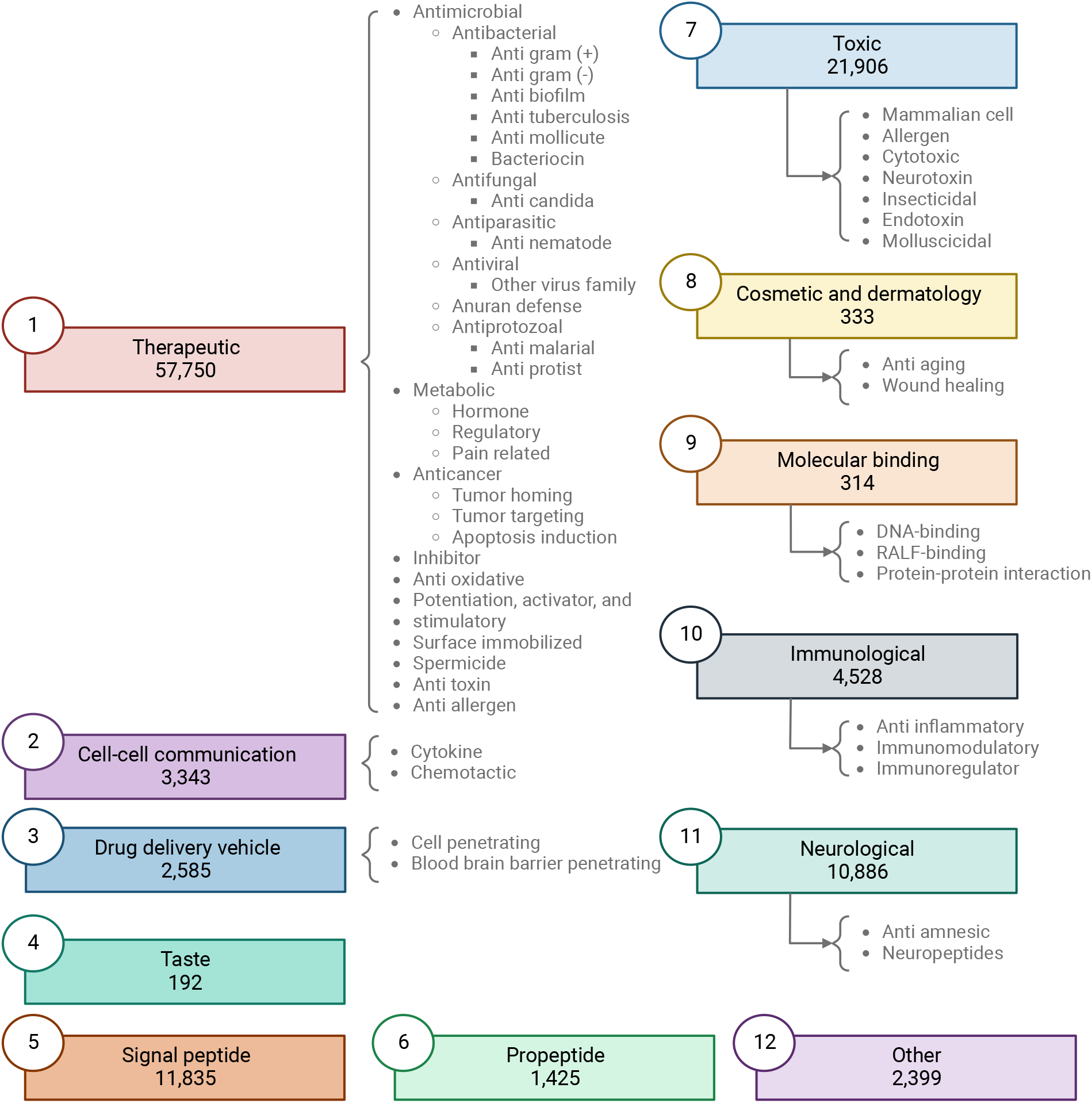
Updated functional biological activity tree implemented in Peptipedia v2.0. In this work, we have updated the functional biological activity tree to include new activities related to taste, molecular binding, and cosmetics and dermatology. Additionally, various antiviral activities have been added, specifically targeting virus families and different mechanisms, with a particular focus on HIV. We have also incorporated peptides associated with therapeutic functions, such as inhibitors, anti-toxins, antiallergens, and spermicides. Peptides with toxic effects have also been included, constituting a significant part of the newly reported peptides in Peptipedia v2.0. Lastly, the category of peptides classified as *“other”* has been updated to encompass diverse biological activities unrelated to the main proposed activities, such as participation in photosynthesis and anti-barnacle properties.

In this new version of the biological activity tree, the activities are organized into 11 primary biological activities at the first level of the tree. Besides, the updated tree has incorporated an additional activity called *“other”*. While maintaining activities associated with therapeutic and immunological effects, this version introduces additional biological activities such as peptides inducing toxic effects, taste peptides, and those with cosmetic and dermatological activities (See Table1 and section S3 of Supplementary Materials for more details).

In Peptipedia v2.0, only 2,399 peptides were categorized as *“other*.*”* These peptides exhibit biological activity but cannot be classified within the biological activities described in the implemented tree. Additionally, only 1,034 peptides—approximately 1% of those registered in this updated version—lack reported activities. This highlights substantial improvements in peptide description and functional understanding compared to the first version reported in Peptipedia (Quiroz et al., 2021).

The updated biological activity tree aims to simplify peptide classification by minimizing ambiguity and facilitating more explicit sequence reporting. Integrating more data sources has resulted in a significant increase in sequences with reported biological activities. Peptipedia v2.0 now incorporates over 100,000 sequences with reported biological activities, representing a surge of over 40% from its initial release (Quiroz et al., 2021). This positions Peptipedia v2.0 not only as the largest repository of peptide sequences but also as the most extensive collection of peptides with therapeutic and antimicrobial activities, exceeding traditional databases such as Erop Moskow (Zamyatnin, 2006), LAMP2 (Ye et al., 2020), DBAMP (Jhong et al., 2021), DBAASP (Pirtskhalava et al., 2020b), and SATPDB (Singh et al., 2015).

### 3.2 Peptipedia incorporates an autonomous updating pipeline

The significant increase in records poses a challenge in maintaining updates, integrating new sequences, and ensuring data integrity. ETL systems (Extract, Transform, and Load) have been designed and implemented to address these database maintenance challenges.

Figure 3 provides an overview of the implemented ETL process for processing a data source. The data source is initially used to extract pertinent information, including peptide sequences, function-associated keywords, literature references, patents, and pharmacological and physicochemical properties. Additionally, metadata is collected and saved for upload into Peptipedia v2.0.

**Figure 3:**
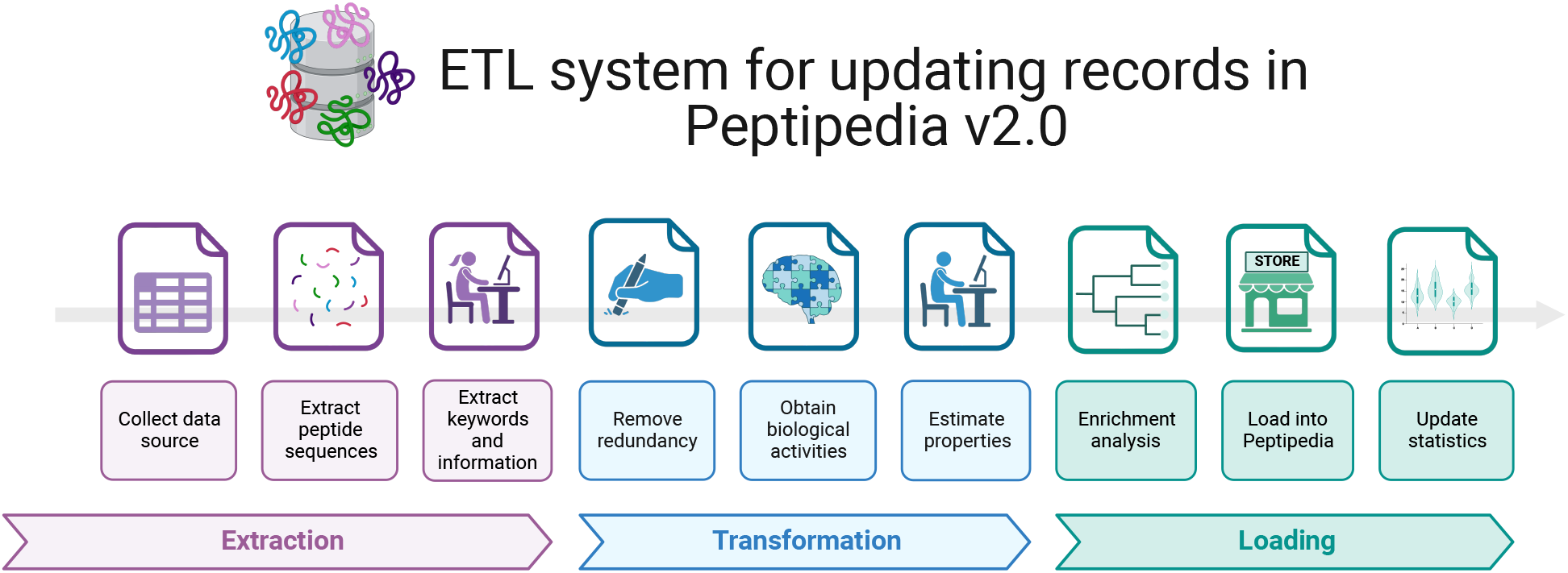
ETL implemented for processing a data source and loading the collected information into Peptipedia DB. The ETL (Extract, Transform, and Load) implemented in Peptipedia facilitates the process of a new data source to extract and load the information into Peptipedia DB. First, the different information is extracted from the data source, including the peptide sequence, references, keywords, and relevant properties described in the data source. Then, two transformation processes are applied to collect raw data to facilitate the characterization of the peptide sequences and the identification of the functional biological activity. Also, an enrichment analysis is run to obtain gene ontology terms and functional domains. Finally, the load process implies inserting or updating the records in the Peptipedia DB, a statistical analysis, and a summary process of the ETL execution.

After data collection and extraction of relevant information, a redundancy evaluation is conducted to remove duplicate elements. The data is then processed to determine biological activity using the functional biological activity tree implemented in Peptipedia v2.0. Peptides are classified as either canonical or non-canonical based on their sequences. For peptides categorized as canonical, physicochemical and thermodynamical properties are estimated using the ModLAMP library (Müller et al., 2017).

Next, an enrichment analysis, including Gene Ontology prediction and functional domain analysis through the Pfam tool, is applied to all canonical peptides.

Once the peptide sequences are processed, the loading process begins, and the peptide sequences are inserted or updated in Peptipedia v2.0. Subsequently, the metadata, statistics, and summary records in Peptipedia v2.0 are updated, completing the process.

An extraction process was developed for each data source to update Peptipedia v2.0. This implementation is critical due to the unique rules and strategies governing the storage and deployment of information within each data source.

Working with each data source individually simplifies the update process, facilitating individualized execution of record updates within Peptipedia v2.0. This flexibility enables tailored configurations for updates based on the update periods stipulated by the data sources. For instance, databases such as UniProt or PDB undergo monthly updates, whereas databases like SATPdb receive annual updates. Conversely, data sources requiring regular updates are included in the Peptipedia v2.0 update process.

### 3.3 Improving functionalities and implementing new bioinformatics tools

Peptipedia v2.0 introduces different tools to facilitate the study of peptide sequences. Firstly, enrichment analysis methods have been integrated, leveraging the predictive capabilities of Gene Ontology terms (Aleksander et al., 2023) and protein functional domains through the Pfam tool (Mistry et al., 2020). Secondly, the prediction of secondary structures and essential structural properties, such as solvent accessibility, is simplified by integrating the RaptorX-Property tool (Wang et al., 2016).

Furthermore, new tools and methods have been implemented to facilitate the application of machine learning algorithms to study peptide sequences. Different numerical representation strategies have been incorporated to simplify the preprocessing of peptide sequences. These strategies encompass coding amino acids based on their physicochemical properties and utilizing learning representations derived from pre-trained models(Medina-Ortiz et al., 2022; Dallago et al., 2021).

The updated machine learning tool now effectively trains predictive models using diverse supervised learning algorithms, including the Gaussian Process, ensemble methods, support vector machines, and nearest neighbour algorithms. Besides, the metrics used to evaluate the training process’s performance have been refined, encompassing performance metrics like Precision, recall and F1-score for classification tasks and Root Mean Squared Error (RMSE) for regression tasks. Both models trained, dataset processed, and summary training process could be downloaded to facilitate using the generated model in a local environment.

Moreover, Peptipedia v2.0 offers pattern recognition capabilities through clustering strategies based on unsupervised learning algorithms, sequence similarity networks, and community detection methods. Peptipedia v2.0 facilitates evaluating the generated groups by applying performance metrics like the Calinski-Harabasz index and the Silhouette coefficients. These algorithms generate cohesive groups characterized using statistical analyses and enrichment tools.

Finally, Peptipedia v2.0 has updated bioinformatic tools for sequence characterization. These include physicochemical property analyses, sequence alignments against the Peptipedia database, and the integration of multiple sequence alignments and statistical description methodologies for groups of protein sequences, which have been implemented to support the study of peptide sequences.

### 3.4 New classification models and predicting non-annotated peptide sequences

The enhancement of the biological activity tree and the expansion of peptide sequences necessitated updating the binary classification models initially developed for Peptipedia. Following the sequence-based approach detailed in the Methods section, this work used ninety-eight biological activities to train binary classification models.

We evaluated 90 combinations of numerical representation strategies and supervised learning algorithms for each biological activity, resulting in over 8,800 trained models. On average, the specificity is 0.807 ± 0.084, and the sensitivity is 0.813 ± 0.082 for the evaluated models (see Section S4 of Supplementary Materials for more details). Algorithms like ExtraTrees and Random Forest demonstrated the highest sensitivity and specificity, with values exceeding 0.83 *±* 0.06. In contrast, Decision Tree-based models showed the lowest performance, with sensitivity and specificity values below 0.74 ± 0.08. Regarding numerical representation strategies, models trained using the pre-trained models ProTrans t5 xlu50 and ProTrans t5 Uniref exhibited the highest performance, with sensitivity and specificity values above 0.82 ± 0.08. Conversely, the ProTrans t5 XLNET model showed the lowest performance, with sensitivity and specificity values below 0.78 ±0.08 (see Section S4 of Supplementary Materials for more details).

The most effective combinations of numerical representation strategies and supervised learning algorithms involved ProTrans t5 Uniref or ProTrans t5 xlu50 with ExtraTrees, achieving average sensitivity and specificity values exceeding 0.85 *±* 0.06. ProTrans t5 XLNET with Decision Tree was the least effective combination, with average sensitivity and specificity values below 0.72 *±* 0.09.

Based on the sensitivity and specificity performance analysis and the criteria described in the methodology section, 98 binary classification models were selected from the evaluated combinations. The chosen models achieved average sensitivities and specificities of 0.877 ± 0.053 and 0.873 ±0.054, respectively. Combinations such as ProTrans t5 Uniref with ExtraTrees, ProTrans t5 Uniref with Gaussian Process, and ProTrans t5 xlu50 with ExtraTrees were the most frequently selected, representing 25% of all evaluated activities. In contrast, combinations like Esm1B with Adaboost, Esm1B with Random Forest, and ProTrans t5 XLNET with XGBoost were the least frequently selected, representing only 0.9% of the total explored activities (see Section 4 of Supplementary Materials for more details).

A ranking of the best models (see Table 2) and the models with the lowest performance (see Table 3) based on the MCC metric was generated. The top models were associated with various target virus activities, including Anti-human parainfluenza virus and Anti-Andes virus, different virus families like *Bunyaviridae* and *Hantaviridae*, and activities such as DNA-Binding, Neurotoxin, and Transit. In contrast, the models with the lowest MCC values were related to activities such as Cosmetic and Dermatology, Blood-Brain Barrier Penetration, and Anticancer. However, despite being classified as the lowest-performing models, their MCC values were higher than 0.5, and they achieved precision averages over 75%, demonstrating high generalization and robustness of the implemented strategies.

**Table 1:** Summary of activities classified in the first level of the updated functional biological tree.

| Biological activity | Definition | Records | Data sources |
| --- | --- | --- | --- |
| Therapeutic | Therapeutic peptides are molecules, whether designed or natural, with medical applications that aim to treat or alleviate diseases by exerting various bodily functions. | 57,750 | 51 |
| Cell-cell communication | Peptides involved in cell-to-cell communication, which can influence cell signaling and cell-mediated immune responses. | 3,343 | 6 |
| Drug delivery vehicle | These peptides function as carriers that transport and deliver drugs in a specific manner to target cells or tissues. They can target specific receptors on cells to release their therapeutic cargo. | 2,858 | 11 |
| Taste | A short chain of amino acids that elicits a specific taste sensation when it interacts with taste receptors on the tongue. Taste peptides can evoke a range of tastes, including sweet, bitter, salty, sour, and umami. | 192 | 2 |
| Signal peptide | Peptides that direct proteins to specific cellular locations used in protein secretion and intracellular transport processes, which may affect cellular communication and protein-mediated immune responses | 11,835 | 9 |
| Propeptide | Inactive peptides that are converted to active peptides by processing, act as a precursor to a protein and are removed during processing, which can influence biological activity and protein regulation. | 1,424 | 2 |
| Toxic | Peptides that possess harmful properties and can induce toxic effects in biological systems. These peptides may interfere with normal cellular functions, alter metabolic processes or cause damage to cells and tissues. | 21,906 | 20 |
| Cosmetic and dermatology | Small-chain amino acid peptides are used in skincare and cosmetic products. These peptides are designed to address various skin-related concerns such as wrinkles, fine lines, loss of elasticity and other signs of skin ageing. | 333 | 5 |
| Molecular binding | Peptides that bind specifically to molecules, such as proteins, RNA, or DNA, which may influence the regulation of biological processes and immune responses. | 314 | 3 |
| Immunological | Peptides have properties that influence the body’s immune response and interact with components of the immune system. They can modulate the function of immune cells and have a role in regulating and modulation of the immune response. | 4258 | 11 |
| Neurological | Essential peptides in nervous system signalling, influencing cellular communication and functions such as memory and neuronal regeneration. | 10,886 | 5 |
| Other | Peptides with other activities non-generalizable with activities in the first level of the tree, including activities like reductant, pore-forming, or ribonucleoprotein | 2,399 | 10 |

**Table 2:** Best models based on MCC performances for binary classification of functional biological activities.

| Activity | Algorithm | Encoder | MCC | Accuracy | F1-score | Precision | Recall |
| --- | --- | --- | --- | --- | --- | --- | --- |
| Anti human parainfluenza virus | RandomForest | Prottrans t5bdf | 0.999 | 0.999 | 0.999 | 0.999 | 0.999 |
| *Anti bunyaviridae* | XGB | Prottrans t5bdf | 0.974 | 0.986 | 0.986 | 0.987 | 0.986 |
| Anti sin nombre virus | RandomForest | Prottrans t5bdf | 0.974 | 0.986 | 0.986 | 0.987 | 0.986 |
| DNA-binding | Bagging | Prottrans t5 uniref | 0.972 | 0.986 | 0.986 | 0.986 | 0.986 |
| Gluten immunogenic and celiac toxic | XGB | Prottrans xlnet | 0.970 | 0.985 | 0.985 | 0.985 | 0.985 |
| *Anti hantaviridae* | AdaBoost | Prottrans xlnet | 0.962 | 0.980 | 0.980 | 0.981 | 0.980 |
| Anti andes virus | RandomForest | Prottrans t5bdf | 0.961 | 0.980 | 0.980 | 0.981 | 0.980 |
| Neurotoxin | GradientBoosting | Esm1b | 0.948 | 0.974 | 0.974 | 0.974 | 0.974 |
| Transit | KNeighbors | Prottrans t5 xlu50 | 0.943 | 0.971 | 0.971 | 0.972 | 0.971 |
| Cytokine | GaussianProcess | Prottrans t5 uniref | 0.910 | 0.955 | 0.955 | 0.955 | 0.955 |

**Table 3:** Models with the lowest MCC performances for binary classification of functional biological activities.

| Activity | Algorithm | Encoder | MCC | Accuracy | F1-score | Precision | Recall |
| --- | --- | --- | --- | --- | --- | --- | --- |
| Anticancer | KNeighbors | Prottrans t5 uniref | 0.607 | 0.800 | 0.799 | 0.806 | 0.800 |
| *Anti retroviridae* | GaussianProcess | Prottrans t5 xlu50 | 0.604 | 0.802 | 0.802 | 0.802 | 0.802 |
| *Anti methicillin-resistant S. aureus* | Bagging | Esm1b | 0.602 | 0.806 | 0.806 | 0.806 | 0.806 |
| Cytotoxic | GaussianProcess | Prottrans t5 uniref | 0.599 | 0.798 | 0.797 | 0.802 | 0.798 |
| Blood brain barrier penetrating | GradientBoosting | Prottrans t5 uniref | 0.591 | 0.796 | 0.796 | 0.796 | 0.796 |
| Anti angiogenic | ExtraTrees | Prottrans t5 uniref | 0.579 | 0.789 | 0.788 | 0.790 | 0.789 |
| Anti biofilm | ExtraTrees | Esm1b | 0.578 | 0.788 | 0.788 | 0.789 | 0.788 |
| Anti influenza virus | ExtraTrees | Prottrans xlnet | 0.576 | 0.790 | 0.789 | 0.790 | 0.790 |
| Cosmetic and dermatology | RandomForest | Prottrans t5bdf | 0.576 | 0.785 | 0.784 | 0.790 | 0.785 |
| Anti candida | ExtraTrees | Prottrans t5 xlu50 | 0.505 | 0.753 | 0.753 | 0.754 | 0.753 |

Finally, the trained classification models evaluated 3.8 million sequences without reported biological activity incorporated into Peptipedia v2.0. Table 4 summarizes the classification results for the biological activities at the first level of the updated biological activity tree. Except for peptides with Molecular Binding activity, all biological activities saw an increase in records by more than 100%. Notably, categories such as therapeutics, signal peptides, propeptides, immunological, and neurological activities increased by more than 600,000 peptides. This suggests a tendency for immunological activity and propeptides among previously uncharacterized peptides. Additionally, more than 800,000 peptides showed potential therapeutic activity, indicating they could serve as alternatives to traditional drug-based compounds. However, further evaluation of these peptides is necessary, including toxicity and immunogenicity assessments and structural analyses to understand their properties compared to previously validated therapeutic peptides (Goles et al., 2024).

**Table 4:**
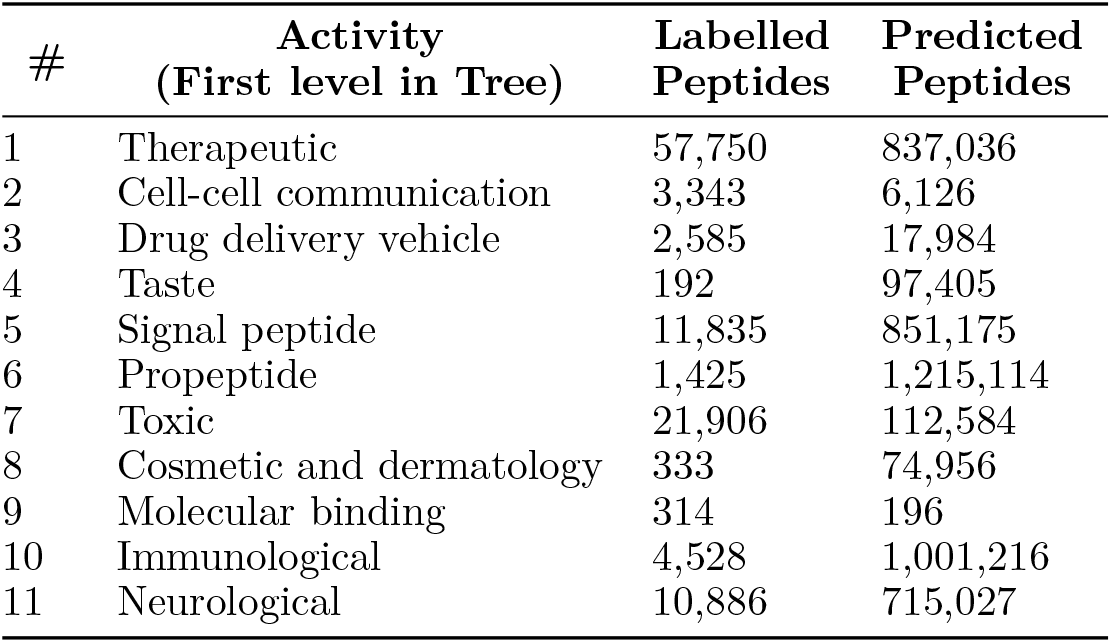
Summary classified unknown peptides used the trained classification models.

### 3.5 A new and modern face for Peptipedia web platform

In this new version of Peptipedia, a visual restructuring of the web platform was designed and implemented. Using web development technologies based on React, a new front-end was developed. The new Peptipedia platform separates the tools for visualizing the database and their respective data searching and downloading options. The separation of tools and databases allows for optimizing response times for each user-generated action (See Figure 4 for a schematic representation and section S5 of Supplementary Materials for more details).

**Figure 4:**
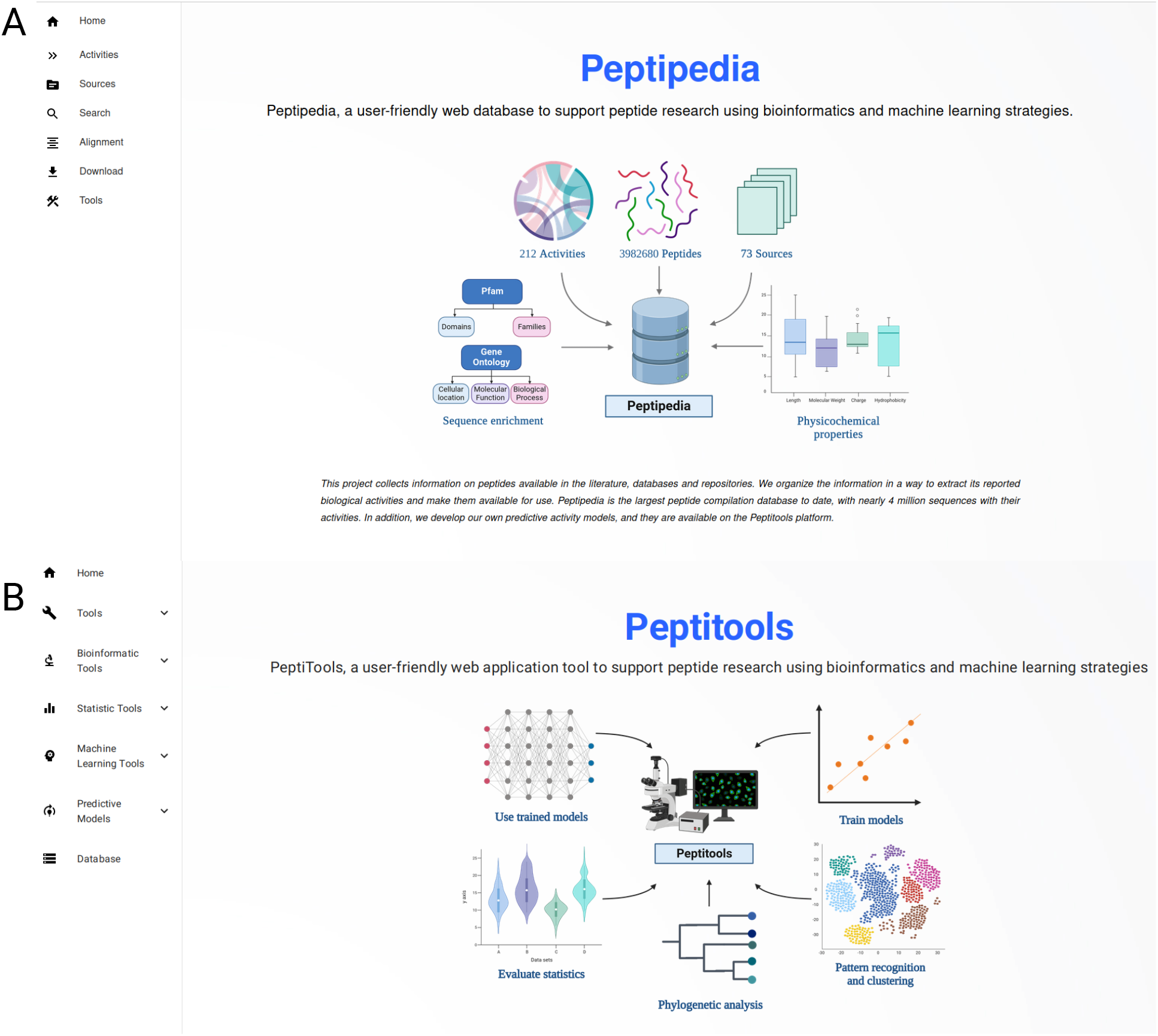
Homepage for Peptipedia DB and Peptitools, the two main services implemented into Peptipedia. **A** Homepage generated to access on PeptipediaDB. This homepage has i) the full menu with all functionalities available on the system, ii) a schematic representation of the implemented workflow to process collected data from data sources, iii) a statistical description of register information, and iv) relevant information on the project. **B** Homepage generated to access on PeptiTools. This homepage includes a schematic representation of implemented tools and a full menu to access a desirable tool.

Upon accessing the Peptipedia web platform via its access link, the user will first see the Peptipedia homepage (See Figure 4 A). This homepage features information about the database, a statistical summary of the information recorded in Peptipedia, a diagram illustrating how data is collected and processed, and information describing the workgroup and the project (See more details in section S5 of Supplementary Materials for more information).

The Peptipedia menu includes the main accesses to the platform, including activity analysis, data sources, direct download systems, and the search engine. The latest version of the sequence search engine in Peptipedia facilitates the application of different filters to customize the type of information identification. Query results have also been optimized through the generation of materialized views. The visualization of the results has been updated to present the characteristics of the identified sequences in a more user-friendly manner, incorporating all existing information in the platform (see Section S5 of Supplementary Material for more details).

From the perspective of the computational tools implemented in Peptipedia, a new platform called PeptiTools has been developed (See Figure 4 B). This platform contains all the tools included in the system, including bioinformatics analysis and enrichment tools, sequence characterization, predictions of biological activities via classification model binaries trained based on the biological activities updated in this new version of Peptipedia, machine learning application methods for building predictive models, and pattern identification methods based on peptide sequence clustering techniques and unsupervised learning algorithms.

Among the new tools incorporated in this latest version of Peptipedia, machine learning methods play an essential role in predicting and studying unknown peptide sequences and training predictive models or pattern recognition.

Only amino acid sequences are required when using classification models of biological activities, and the system will automatically predict the selected biological activities. To do this, the system collects the sequences, numerically represents them according to the pre-trained model corresponding to the activity of interest, applies the model, and delivers the responses. A configurable probability threshold parameter is incorporated to customize predictions and provide greater decision control to the user.

The training of predictive models can be carried out using the corresponding tool implemented in PeptiTools (see Section S5 of Supplementary Materials for more details). These models are based on numerical rep-resentation of sequences, so they do not employ structural techniques or feature engineering, only allowing the application of pre-trained models and encodings based on physicochemical properties. Besides, different configurations for preprocessing and training models are available, including i) selecting the algorithm to train the model, ii) validation strategies, and iii) standardization and dimensionality reduction applications. Once the model is trained, the results are displayed on the platform, which varies depending on the type of response used to train the model, indicating specific information such as scatter plots for regression models and sensitivity-specificity analysis with confusion matrices for classification models.

When applying clustering strategies for pattern recognition, the tool requires input sequences, the selection of a numerical representation strategy, and the choice of methodology. The tool processes the entered configuration, applies the approach, and generates the results, displaying the characterized groups and a summary of the process (See section S5 of Supplementary materials for more details).

## 4 Conclusions

This work introduces Peptipedia v2.0, a significant update to our previously reported peptide database. Peptipedia v2.0 integrates data from over 70 sources, compiling over 100,000 peptides with reported biological activity and over 3.8 million peptide sequences without reported biological activity. The functional biological activity tree has been updated, increasing the number of biological activities and refining the hierarchical structure to facilitate the efficient study of peptide functions.

In this new version of Peptipedia, all registered peptides are characterized using enrichment analysis approaches, integrating gene ontology term predictions, secondary structure evaluation, and functional domain analysis. Additionally, various physicochemical and thermodynamic properties are estimated for each peptide sequence, enhancing the knowledge base of Peptipedia v2.0. The database also includes patents, references, and pharmacological properties related to the peptide sequences.

The classification models have been enhanced with the updated functional biological activity tree. A combination of numerical representation strategies based on embedding approaches through pre-trained models and classical supervised learning algorithms was used to train the classification models, achieving an average sensitivity and specificity of 87% across more than 95 evaluated models. These models were utilized to identify potential therapeutic peptides and predict the functional biological activity of the 3.8 million previously uncharacterized peptide sequences.

Furthermore, Peptipedia v2.0 incorporates new services and tools, including enrichment analysis, statistical evaluation, and machine learning-based strategies, to facilitate the study of peptide sequences. By updating the database and developing these tools, Peptipedia v2.0 positions itself as one of the most comprehensive public repositories for peptide research, playing a crucial role in various research areas.

## Supporting information

Supplentary Information of Peptipedia V2.0 work.

## 6 Author contributions statement

GC-M, and DM-O: conceptualization. DM-O, RU-P and GC-M: methodology. DM-O, AD, and MAN: validation. GC-M, LS-V, VG, AD, and DA-S: investigation. DM-O, AD, JS-Y, GC-M, MAN, RU-P, ONNFC, and MDD: writing, review, and editing. DM-O and RU-P: supervision and funding resources. DM-O and RU-P: project administration.

## 7 Acknowledgments

DM-O acknowledges ANID for the project “SUBVENCIóN A INSTALACIóN EN LA ACADEMIA CONVOCATORIA AñO 2022”, Folio 85220004. DM-O, RU-P, and AD gratefully acknowledge support from the Centre for Biotechnology and Bioengineering - CeBiB (PIA project FB0001, Conicyt, Chile), and MAN acknowledges ANID for grants Anillo ATE220016. MAN and RU-P acknowledge ANID for grant Fondecyt 1230298. MDD acknowledges funding by the Deutsche Forschungsgemeinschaft (DFG, German Research Foundation) - SPP2363. PEACCEL was supported through a research program partially cofunded by the European Union (UE) and Region Reunion (FEDER).

## 8 Data Availability Statement

Source code and datasets are available on the GitHub repository: i) Peptipedia v2.0 database https://github.com/ProteinEngineering-PESB2/Peptipedia and ii)Peptipedia v2.0 tools: https://github.com/ProteinEngineering-PESB2/Peptitools. The ETL process scripts are located in https://github.com/ProteinEngineering-PESB2/PeptipediaParser. The user-friendly web platform is publically accessible through https://app.peptipedia.cl/ for non-commercial uses, licensed under a Creative Commons CC BY-NC-ND 4.0 license. The Peptipedia v2.0 database is available for non-commercial use and is licensed under an ODbl license.

## References

(2023). Uniprot: the universal protein knowledgebase in 2023. Nucleic Acids Research, 51(D1):D523–D531.

Aleksander, S. A., Balhoff, J., Carbon, S., Cherry, J. M., Drabkin, H. J., Ebert, D., Feuermann, M., Gaudet, P., Harris, N. L., et al. (2023). The gene ontology knowledgebase in 2023. Genetics, 224(1):iyad031.

Apostolopoulos, V., Bojarska, J., Chai, T.-T., Elnagdy, S., Kaczmarek, K., Matsoukas, J., New, R., Parang, K., Lopez, O. P., Parhiz, H., et al. (2021). A global review on short peptides: Frontiers and perspectives. Molecules, 26:430.

Ashburner, M., Ball, C. A., Blake, J. A., Botstein, D., Butler, H., Cherry, J. M., Davis, A. P., Dolinski, K., Dwight, S. S., Eppig, J. T., et al. (2000). Gene ontology: tool for the unification of biology. Nature genetics, 25(1):25–29.

Berman, H. M., Westbrook, J., Feng, Z., Gilliland, G., Bhat, T. N., Weissig, H., Shindyalov, I. N., and Bourne, P. E. (2000a). The protein data bank. Nucleic acids research, 28(1):235–242.

Berman, H. M., Westbrook, J., Feng, Z., Gilliland, G., Bhat, T. N., Weissig, H., Shindyalov, I. N., and Bourne, P. E. (2000b). The Protein Data Bank. Nucleic Acids Research, 28(1):235–242.

Biswas, S., Khimulya, G., Alley, E. C., Esvelt, K. M., and Church, G. M. (2021). Low-n protein engineering with data-efficient deep learning. Nature methods, 18(4):389–396.

Dallago, C., Schütze, K., Heinzinger, M., Olenyi, T., Littmann, M., Lu, A. X., Yang, K. K., Min, S., Yoon, S., Morton, J. T., and Rost, B. (2021). Learned embeddings from deep learning to visualize and predict protein sets. Current Protocols, 1(5):e113.

Goeddel, D. V., Kleid, D. G., Bolivar, F., Heyneker, H. L., Yansura, D. G., Crea, R., Hirose, T., Kraszewski, A., Itakura, K., and Riggs, A. D. (1979). Expression in escherichia coli of chemically synthesized genes for human insulin. Proceedings of the National Academy of Sciences, 76(1):106–110.

Goles, M., Daza, A., Cabas-Mora, G., Sarmiento-Varón, L., Sepúlveda-Yañez, J., Anvari-Kazemabad, H., Davari, M. D., Uribe-Paredes, R., Olivera-Nappa, A., Navarrete, M. A., and Medina-Ortiz, D. (2024). Peptide-based drug discovery through artificial intelligence: towards an autonomous design of therapeutic peptides. Briefings in Bioinformatics, 25(4):bbae275.

GVR Report Cover (2023). Peptide therapeutics market analysis, 2018-2030 — base year - 2022. Electronic (PDF). Report ID: 978-1-68038-179-5, Number of Pages: 110, Historical Range: 2018 - 2021, Industry: Healthcare.

Hammami, R., Zouhir, A., Le Lay, C., Hamida, J. B., and Fliss, I. (2010). Bactibase second release: a database and tool platform for bacteriocin characterization. Bmc Microbiology, 10(1):1–5.

Hamp, T., Kassner, R., Seemayer, S., Vicedo, E., Schaefer, C., Achten, D., Auer, F., Boehm, A., Braun, T., Hecht, M., Heron, M., Hönigschmid, P., Hopf, T. A., Kaufmann, S., Kiening, M., Krompass, D., Landerer, C., Mahlich, Y., Roos, M., and Rost, B. (2013). Homology-based inference sets the bar high for protein function prediction. BMC Bioinformatics, 14(S3).

Henninot, A., Collins, J. C., and Nuss, J. M. (2018). The current state of peptide drug discovery: back to the future? Journal of medicinal chemistry, 61(4):1382–1414.

Jeffery, C. J. (1999). Moonlighting proteins. Trends in biochemical sciences, 24(1):8–11.

Jhong, J.-H., Yao, L., Pang, Y., Li, Z., Chung, C.-R., Wang, R., Li, S., Li, W., Luo, M., Ma, R., Huang, Y., Zhu, X., Zhang, J., Feng, H., Cheng, Q., Wang, C., Xi, K., Wu, L.-C., Chang, T.-H., Horng, J.-T., Zhu, L., Chiang, Y.-C., Wang, Z., and Lee, T.-Y. (2021). dbAMP 2.0: updated resource for antimicrobial peptides with an enhanced scanning method for genomic and proteomic data. Nucleic Acids Research, 50(D1):D460–D470.

Jumper, J., Evans, R., Pritzel, A., Green, T., Figurnov, M., Ronneberger, O., Tunyasuvunakool, K., Bates, R., Žídek, A., Potapenko, A., et al. (2021). Highly accurate protein structure prediction with alphafold. Nature, 596(7873):583–589.

Kang, X., Dong, F., Shi, C., Liu, S., Sun, J., Chen, J., Li, H., Xu, H., Lao, X., and Zheng, H. (2019). Dramp 2.0, an updated data repository of antimicrobial peptides. Scientific data, 6(1):1–10.

Khan, M. M., Filipczak, N., and Torchilin, V. P. (2021). Cell penetrating peptides: A versatile vector for co-delivery of drug and genes in cancer. Journal of Controlled Release, 330:1220–1228.

Lau, J. L. and Dunn, M. K. (2018). Therapeutic peptides: Historical perspectives, current development trends, and future directions. Bioorganic & medicinal chemistry, 26(10):2700–2707.

Lee, A. C.-L., Harris, J. L., Khanna, K. K., and Hong, J.-H. (2019). A comprehensive review on current advances in peptide drug development and design. International journal of molecular sciences, 20(10):2383.

Lien, S. and Lowman, H. B. (2003). Therapeutic peptides. Trends in biotechnology, 21(12):556–562.

Medina-Ortiz, D., Contreras, S., Amado-Hinojosa, J., Torres-Almonacid, J., Asenjo, J. A., Navarrete, M., and Olivera-Nappa, Á. (2020a). Combination of digital signal processing and assembled predictive models facilitates the rational design of proteins. arXiv preprint 2010.03516.

Medina-Ortiz, D., Contreras, S., Amado-Hinojosa, J., Torres-Almonacid, J., Asenjo, J. A., Navarrete, M., and Olivera-Nappa, Á. (2022). Generalized property-based encoders and digital signal processing facilitate predictive tasks in protein engineering. Frontiers in Molecular Biosciences, 9.

Medina-Ortiz, D., Contreras, S., Quiroz, C., Asenjo, J. A., and Olivera-Nappa, Á. (2020b). Dmakit: A user-friendly web platform for bringing state-of-the-art data analysis techniques to non-specific users. Information Systems, page 101557.

Medina-Ortiz Sr, D., Cabas-Mora Sr, G., Moya-Barria Sr, I., Soto-Garcia, N., and Uribe-Paredes Sr, R. (2024). Rudeus, a machine learning classification system to study dna-binding proteins. bioRxiv, pages 2024–02.

Mistry, J., Chuguransky, S., Williams, L., Qureshi, M., Salazar, G. A., Sonnhammer, E. L., Tosatto, S. C., Paladin, L., Raj, S., Richardson, L. J., et al. (2021). Pfam: The protein families database in 2021. Nucleic acids research, 49(D1):D412–D419.

Mistry, J., Chuguransky, S., Williams, L., Qureshi, M., Salazar, G. A., Sonnhammer, E. L. L., Tosatto, S. C. E., Paladin, L., Raj, S., Richardson, L. J., Finn, R. D., and Bateman, A. (2020). Pfam: The protein families database in 2021. Nucleic Acids Research, 49(D1):D412–D419.

Muttenthaler, M., King, G. F., Adams, D. J., and Alewood, P. F. (2021). Trends in peptide drug discovery. Nature reviews Drug discovery, 20(4):309–325.

Müller, A. T., Gabernet, G., Hiss, J. A., and Schneider, G. (2017). modlAMP: Python for antimicrobial peptides. Bioinformatics, 33(17):2753–2755.

Pirtskhalava, M., Amstrong, A. A., Grigolava, M., Chubinidze, M., Alimbarashvili, E., Vishnepolsky, B., Gabrielian, A., Rosenthal, A., Hurt, D. E., and Tartakovsky, M. (2020a). Dbaasp v3: database of antimicrobial/cytotoxic activity and structure of peptides as a resource for development of new therapeutics. Nucleic Acids Research.

Pirtskhalava, M., Amstrong, A. A., Grigolava, M., Chubinidze, M., Alimbarashvili, E., Vishnepolsky, B., Gabrielian, A., Rosenthal, A., Hurt, D. E., and Tartakovsky, M. (2020b). DBAASP v3: database of antimicrobial/cytotoxic activity and structure of peptides as a resource for development of new therapeutics. Nucleic Acids Research, 49(D1):D288–D297.

Quiroz, C., Saavedra, Y. B., Armijo-Galdames, B., Amado-Hinojosa, J., Olivera-Nappa, Á., Sanchez-Daza, A., and Medina-Ortiz, D. (2021). Peptipedia: a user-friendly web application and a comprehensive database for peptide research supported by machine learning approach. Database, 2021:baab055.

Ramaprasad, A. S. E., Singh, S., Venkatesan, S., et al. (2015). Antiangiopred: a server for prediction of anti-angiogenic peptides. PloS one, 10(9):e0136990.

Sims, E. K., Carr, A. L., Oram, R. A., DiMeglio, L. A., and Evans-Molina, C. (2021). 100 years of insulin: celebrating the past, present and future of diabetes therapy. Nature medicine, 27(7):1154–1164.

Singh, N. and Bhalla, N. (2020). Moonlighting proteins. Annual review of genetics, 54:265–285.

Singh, S., Chaudhary, K., Dhanda, S. K., Bhalla, S., Usmani, S. S., Gautam, A., Tuknait, A., Agrawal, P., Mathur, D., and Raghava, G. P. (2015). SATPdb: a database of structurally annotated therapeutic peptides. Nucleic Acids Research, 44(D1):D1119–D1126.

Singh, S., Chaudhary, K., Dhanda, S. K., Bhalla, S., Usmani, S. S., Gautam, A., Tuknait, A., Agrawal, P., Mathur, D., and Raghava, G. P. (2016). Satpdb: a database of structurally annotated therapeutic peptides. Nucleic acids research, 44(D1):D1119–D1126.

Taylor, S. I. (2000). Rational design of peptide agonists of cell-surface receptors. Trends in Pharmacological Sciences, 21(1):9–10.

Van Dorpe, S., Bronselaer, A., Nielandt, J., Stalmans, S., Wynendaele, E., Audenaert, K., Van De Wiele, C., Burvenich, C., Peremans, K., Hsuchou, H., et al. (2012). Brainpeps: the blood–brain barrier peptide database. Brain Structure and Function, 217(3):687–718.

Wan, F., Kontogiorgos-Heintz, D., and de la Fuente-Nunez, C. (2022). Deep generative models for peptide design. Digital Discovery, 1(3):195–208.

Wang, L., Wang, N., Zhang, W., Cheng, X., Yan, Z., Shao, G., Wang, X., Wang, R., and Fu, C. (2022). Therapeutic peptides: Current applications and future directions. Signal Transduction and Targeted Therapy, 7(1):48.

Wang, S., Li, W., Liu, S., and Xu, J. (2016). RaptorX-Property: a web server for protein structure property prediction. Nucleic Acids Research, 44(W1):W430–W435.

Wynendaele, E., Bronselaer, A., Nielandt, J., D’Hondt, M., Stalmans, S., Bracke, N., Verbeke, F., Van De Wiele, C., De Tre, G., and De Spiegeleer, B. (2013). Quorumpeps database: chemical space, microbial origin and functionality of quorum sensing peptides. Nucleic acids research, 41(D1):D655–D659.

Ye, G., Wu, H., Huang, J., Wang, W., Ge, K., Li, G., Zhong, J., and Huang, Q. (2020). LAMP2: a major update of the database linking antimicrobial peptides. Database, 2020.

Zamyatnin, A. (1991). Erop-moscow: specialized data bank for endogenous regulatory oligopeptides. Protein sequences & data analysis, 4(1):49–52.

Zamyatnin, A. A. (2006). The EROP-moscow oligopeptide database. Nucleic Acids Research, 34(90001):D261–D266.

Zanzoni, A., Ribeiro, D. M., and Brun, C. (2019). Understanding protein multifunctionality: from short linear motifs to cellular functions. Cellular and Molecular Life Sciences, 76:4407–4412.

Zhao, X., Wu, H., Lu, H., Li, G., and Huang, Q. (2013). Lamp: a database linking antimicrobial peptides. PloS one, 8(6):e66557.

