## Supplementary material for "Peptipedia v2.0: A peptide sequence database and user-friendly web platform. A major update": Supplentary Information of Peptipedia V2.0 work.

Supplementary Information

#### S1 Incorporated databases into Peptipedia v2.0

76 sources were processed to collect the information on peptide sequences incorporated in the new version of Peptipedia v2.0. In this version, it was incorporated databases and

Table S1: Databases and sources collected to incorporate in Peptipedia v2.0

| # | Source or database | Collected peptides | Reference | Database or Dataset |
|---|---|---|---|---|
| 1 | Peptide Atlas | 3747783 | [16] | Database |
| 2 | Swiss-Prot | 106766 | [6] | Database |
| 3 | RCSB PDB | 49457 | [9] | Database |
| 4 | Erop Moskow | 25948 | [76] | Database |
| 5 | LAMP2 | 22164 | [73] | Database |
| 6 | DBAMP | 21834 | [26] | Database |
| 7 | Ensemble Amppred | 21548 | [37] | Dataset |
| 8 | DBAASP | 16722 | [51] | Database |
| 9 | SATPDB | 15937 | [60] | Database |
| 10 | PeptideDB | 14444 | [5] | Database |
| 11 | Adam | 8097 | [36] | Database |
| 12 | CAMP | 6695 | [69] | Database |
| 13 | Ampfun | 6499 | [12] | Database |
| 14 | DRAMP | 5551 | [28] | Database |
| 15 | Conoserver | 4885 | [27] | Database |
| 16 | Amplify | 4164 | [38] | Dataset |
| 17 | Biopep | 3907 | [44] | Database |
| 18 | PlantPepDB | 3748 | [15] | Database |
| 19 | PREAIP | 3565 | [31] | Dataset |
| 20 | APD3 | 3485 | [70] | Database |
| 21 | AMPEPPY | 3139 | [35] | Dataset |
| 22 | Cancergram analysis | 2801 | [8] | Dataset |
| 23 | Pep-lab DB | 2746 | [62] | Database |
| 24 | AVPiden | 2662 | [48] | Dataset |
| 25 | Yadamp | 2525 | [50] | Database |
| 26 | Predneurop | 2425 | [7] | Dataset |
| 27 | Toxinpred | 2108 | [24] | Dataset |
| 28 | Dravp | 1835 | [39] | Database |
| 29 | AVPDB | 1817 | [54] | Database |
| 30 | DADP | 1792 | [46] | Database |
| 31 | APIN | 1777 | [61] | Dataset |
| 32 | AMP Scan 2 | 1777 | [68] | Dataset |
| 33 | Arachnoserver | 1723 | [49] | Database |
| 34 | AMP | 1710 | [58] | Dataset |

| | | | | |
|---|---|---|---|---|
| 35 | AHTPDB | 1694 | [33] | Database |
| 36 | Hemolytik | 1530 | [22] | Database |
| 37 | ANTIFP | 1459 | [2] | Dataset |
| 38 | Compare | 1201 | [67] | Database |
| 39 | STRAPEP | 1201 | [71] | Database |
| 40 | CPPsite | 1198 | [30] | Database |
| 41 | BiodadPep | 1136 | [59] | Database |
| 42 | Celiac database | 1040 | [4] | Database |
| 43 | Hemopi | 994 | [11] | Dataset |
| 44 | ANTICP2 | 970 | [1] | Dataset |
| 45 | IACP DRLF | 970 | [41] | Dataset |
| 46 | PepLife | 938 | [42] | Database |
| 47 | HIPDB | 888 | [53] | Database |
| 48 | Ensemble classifier chain model | 863 | [77] | Dataset |
| 49 | Avpic50pred | 760 | [52] | Dataset |
| 50 | Allergen online | 740 | [23] | Database |
| 51 | Tumorhope | 704 | [29] | Database |
| 52 | Ennavia | 668 | [63] | Dataset |
| 53 | FermFooDB | 575 | [10] | Dataset |
| 54 | Cellppd | 568 | [21] | Dataset |
| 55 | Neuropedia | 537 | [32] | Database |
| 56 | ACP DL | 505 | [74] | Dataset |
| 57 | Anticancer peptides cnn | 505 | [3] | Dataset |
| 58 | Signal peptide | 389 | [18] | Database |
| 59 | Quorumpeps | 354 | [72] | Database |
| 60 | AntiTB PDB | 346 | [66] | Database |
| 61 | Deepacp | 341 | [75] | Dataset |
| 62 | Brainpeps | 335 | [17] | Database |
| 63 | Vaxinpad | 304 | [45] | Dataset |
| 64 | Aagingbase | 282 | [55] | Database |
| 65 | Anticp | 275 | [64] | Dataset |
| 66 | B3PRED | 265 | [34] | Dataset |
| 67 | Plifepred | 234 | [43] | Dataset |
| 68 | AcovPepDB | 228 | [78] | Database |
| 69 | QSPPred | 220 | [56] | Dataset |
| 70 | Antiangiopred | 202 | [57] | Dataset |
| 71 | Baamps | 189 | [40] | Database |
| 72 | BBPred | 119 | [13] | Dataset |
| 73 | Kelm cpppred | 96 | [47] | Dataset |
| 74 | Bactibase | 86 | [25] | Database |
| 75 | Cancer ppd | 20 | [65] | Database |
| 76 | Marine algae derived | 16 | [20] | Dataset |

#### S2 Pre-trained models employed to represent numerically the peptide sequences

Seven pre-trained models were utilized to represent peptide sequences numerically. These models were collected and applied, leveraging a bio-embedding tool. Table S2 provides a comprehensive overview of these models, detailing their names, references, and the dimensions of the generated tensors.

Table S2: Summary of pre-trained models employed to represent numerically the peptide sequences

| # | Pre-trained model | Description | Tensor size | Reference |
|---|---|---|---|---|
| 1 | ProTrans t5 UniRef | The ProtTrans UniRef pre-trained model is a deep learning model specifically trained for protein sequence representation and understanding. It is trained on the UniRef50 database, which contains clustered protein sequences to reduce redundancy and improve diversity. | 1024 | [19] |
| 2 | ProTrans t5 xlu50 | ProtT5-XL-UniRef50 is based on the t5-3b model and was pre-trained on a a large corpus of protein sequences in a self-supervised fashion. This means it was pre-trained on the raw protein sequences only, with no humans labelling them in any way (which is why it can be used a lot of publicly available data) with an automatic process to generate inputs and labels from that protein sequences. | 1024 | [19] |
| 3 | ProTrans T5-BDF | ProtT5-XL-BFD is based on the t5-3b model and was trained on a large corpus of protein sequences in a self-supervised fashion. This means it was trained on the raw protein sequences only, with no human labelling them in any way (which is why it can use lots of publicly available data) with an automatic process to generate inputs and labels from that protein sequences. | 1024 | [19] |
| 4 | Esm1b | The ESM-1b (Evolutionary Scale Modeling) pre-trained model is a variant of the ESM model, designed for protein sequence modelling. It is based on self-supervised learning techniques and utilizes a Transformer architecture, similar to those used in natural language processing tasks. | 1280 | [14] |
| 5 | ProTrans XLNet | The ProtTrans XLNet pre-trained model is a variant of the XLNet model customized for protein sequence analysis. XLNet is an extension of the transformer-based architecture, which integrates bidirectional context learning with permutation-based training. Similarly, ProtTrans XLNet leverages these features to learn contextual representations of amino acids in protein sequences. | 1024 | [19] |
| 6 | ProTrans ALBERT | The ProtTrans ALBERT (A Lite BERT) pre-trained model is a variant of the ALBERT model specifically adapted for protein sequence analysis. ALBERT is a lightweight version of the BERT model, designed to reduce computational resources while maintaining performance. Similarly, ProtTrans ALBERT leverages this efficiency to provide effective representations of amino acids in protein sequences. | 4096 | [19] |
| 7 | ProTrans BERT | The ProtTrans BERT (Bidirectional Encoder Representations from Transformers) pre-trained model is a variant of the BERT model specifically tailored for protein sequence analysis. Like its counterpart in natural language processing, ProtTrans BERT utilizes a transformer-based architecture to learn contextual representations of amino acids in protein sequences. | 1024 | [19] |

#### S3 Updating the functional biological tree

Peptipedia v2.0, a new functional biological tree, was implemented. The first level of the functional tree includes 11 biological activities. Compared with the previous version of Peptipedia, different biological activities related to the family of antivirals and target viruses were incorporated. Activities like molecular binding and cosmetic and dermatology were also incorporated. A summary of the functional biological tree with all integrated properties and the number of records by activity is summarized below.

All activities

1. Taste (192)
2. Toxic (21906)

(a) Anti endotoxin (95)
(b) Insecticidal (336)
(c) Anti mammalian cell (9494)
(d) Cytotoxic (1511)
    i. Hemolytic (1341)
(e) Neurotoxin (592)
(f) Allergen (2462)
    i. Gluten immunogenic + celiac toxic (1240)
(g) Molluscicidal (6)

3. Therapeutic (57750)
    (a) Spermicide (26)
    (b) Sperm activating (46)
    (c) Anti allergen (11)
    (d) Antimicrobial (40853)
        i. Antiparasitic (6312)
            - Anti nematode (118)
        ii. Antiviral (5899)
            - Anti adenoviridae (6)
                - Anti adenovirus (6)
            - Anti alloherpesviridae (2)
                - Anti channel catfish virus (2)
            - Anti arenaviridae (5)
                - Anti pichinde virus (2)
                - Anti tacaribe virus (1)
                - Anti lymphocytic choriomeningitis virus (1)
                - Anti junin virus (2)
            - Anti arteriviridae (16)
                - Anti porcine reproductive and respiratory syndrome virus (16)
            - Anti caliciviridae (1)
                - Anti murine norovirus (1)
            - Anti asfarviridae (1)
                - Anti african swine fever virus (1)
            - Anti bunyaviridae (126)
                - Anti sin nombre virus (126)
            - Anti coronaviridae (379)
                - Anti SARS-COV 2 (296)
                - Anti canine coronavirus (9)
                - Anti feline coronavirus (3)
                - Anti porcine respiratory coronavirus (1)
                - Anti transmissible gastroenteritis virus (3)
                - Anti sl-cov-wiv1 (1)
                - Anti SARS-COV (130)
                - Anti murine hepatitis virus (2)
                - Anti MERS-COV (49)
                - Anti human coronavirus (21)

- Anti coronavirus (109)
- Anti filoviridae (680)
  - Anti ebola virus (16)
  - Anti hepatitis C virus (481)
  - Anti dengue virus (74)
  - Anti west nile virus (45)
  - Anti japanese encephalitis virus (25)
  - Anti zika virus (11)
  - Anti mouse hepatitis virus (10)
  - Anti hepatitis E virus (10)
  - Anti hepatitis B virus (57)
  - Anti tembusu virus (2)
  - Anti yellow fever virus (1)
- Anti flaviviridae (12)
  - Anti classical swine fever virus (12)
- Anti hantaviridae (86)
  - Anti puumala virus (1)
- Anti herpesviridae (381)
  - Anti herpes simplex virus (327)
  - Anti human cytomegalovirus (45)
  - Anti marek's disease virus (10)
  - Anti varicella zoster virus (3)
  - Anti bovine herpesvirus type 1 (1)
- Anti iridoviridae (10)
  - Anti frog virus (8)
  - Anti singapore grouper iridovirus (2)
- Anti nimaviridae (3)
  - Anti white spot syndrome virus (3)
- Anti nodaviridae (2)
  - Anti nervous necrosis virus (2)
- Anti hantaviridae (86)
  - Anti andes virus (85)
- Anti orthoherpesviridae (5)
  - Anti kaposi's sarcoma-associated herpes virus (3)
  - Anti epstein-barr virus (2)
- Anti orthomyxoviridae (190)
  - Anti influenza virus (179)
  - Anti H1N1 virus (17)
  - Anti H3N2 virus (8)
  - Anti H5N1 virus (1)
  - Anti avian influenza virus (2)
- Anti papillomaviridae (24)
  - Anti human papiloma virus (24)
- Anti paramyxoviridae (344)
  - Anti respiratory syncytial virus (173)
  - Anti human parainfluenza virus (99)
  - Anti newcastle disease virus (18)

  - Anti metapneumovirus (5)
  - Anti sendai virus (8)
  - Anti nipah virus (1)
  - Anti measles virus (46)
  - Anti human metapneumovirus (9)
  - Anti hendra virus (11)
- Anti parvoviridae (5)
  - Anti minute virus of mice (3)
  - Anti mink enteritis virus (2)
- Anti phenuiviridae (1)
  - Anti rift valley fever virus (1)
- Anti picornaviridae (49)
  - Anti coxsackie virus (17)
  - Anti foot-and-mouth disease virus (5)
  - Anti rhinovirus (1)
  - Anti enterovirus (7)
  - Anti encephalomyocarditis virus (14)
  - Anti duck hepatitis virus (5)
- Anti polyomaviridae (5)
  - Anti bk virus (5)
- Anti poxviridae (60)
  - Anti vaccinia (60)
  - Anti monkeypox virus (1)
  - Anti cowpox virus (1)
- Anti reoviridae (2)
  - Anti simian rotavirus (1)
  - Anti bovine rotavirus (1)
- Anti retroviridae (1545)
  - Anti feline immunodeficiency virus (124)
  - Anti simian immunodeficiency virus (19)
  - Anti human t-lymphotropic virus (5)
  - Anti HIV (1402)
  - Anti murine leukemia virus (1)
  - Anti human t-cell leukaemia virus 1 (7)
  - Anti feline leukemia virus (13)
  - Anti avian sarcoma and leukosis virus- a (6)
  - Anti avian myeloblastosis virus (1)
- Anti rhabdoviridae (46)
  - Anti vesicular stomatistic virus (17)
  - Anti rabies virus (29)
- Anti sedoreoviridae (1)
  - Anti grass carp hemorrhage virus (1)
- Anti togaviridae (1)
  - Anti chikungunya virus (1)

iii. Antibacterial (24377)
- Anti gram (+) (18734)
  - Anti methicillin-resistant s. aureus (395)

        - Anti gram (-) (15453)
        - Anti biofilm (505)
        - Anti tuberculosis (357)
        - Bacteriocin (2)
        - Anti mollicute (39)
    iv. Anuran defense (1806)
    v. Anti fungal (11772)
        - Anti candida (758)
    vi. Antiprotozoal (165)
        - Anti protist (42)
        - Anti malarial (106)

(e) Anti toxin (19)

(f) Anticancer (9724)
    i. Tumor homing (369)
    ii. Tumor targeting (344)
    iii. Apoptosis induction (1)

(g) Metabolic (13487)
    i. Pain related (173)
        - Opioid (165)
            - Opioid-antagonist (9)
            - Opioid-agonist (14)
    ii. Hormone (9307)
        - Pheromone (61)
    iii. Regulatory (4070)
        - Coagulation + vascular (2214)
            - Anti angiogenic (280)
                * Venous (68)
                    · Anti thrombotic (68)
            - Anti hypertensive (1859)
            - Anti hypotensive (32)
        - Glucose (1514)
            - Anti diabetic (1514)
                * Anti diabetic type 1 (685)
                * Anti diabetic type 2 (802)
        - Quorum sensing (354)

(h) Inhibitor (3523)
    i. Enzyme inhibitor (3324)
        - Protease inhibitor (70)
        - Glucosidase inhibitor
        - Amylase inhibitor (40)
        - Angiotensinase inhibitor (1876)
            - Angiotensin-converting enzyme (ace) inhibitors (1871)

(i) Activator (112)
    i. Enzyme activator (62)

(j) Stimulator (112)

(k) Potentiator (222)
(l) Anti oxidative (1117)
(m) Surface immobilized (46)
(n) Antiatherosclerotic (4)
(o) Antiosteoporosis (1)

4. Neurological (71)
   (a) Anti amnesic (65)
5. Immunological (4528)
   (a) Immunomodulatory (433)
   (b) Immunoregulator (158)
   (c) Anti inflamatory (3902)
6. Molecular binding (314)
   (a) Protein-protein interaction (9)
   (b) DNA-binding (119)
   (c) Ralf-binding (41)
7. Propeptide (1425)
8. Cosmetic and dermatology (333)
   (a) Wound healing (45)
   (b) Anti aging (282)
9. Other (7406)
10. Signal peptide (22650)
   (a) Neuropeptide (10815)
   (b) Transit (7523)
11. Drug delivery vehicle (2585)
   (a) Cell penetrating (1338)
   (b) Blood brain barrier penetrating (561)
12. Cell cell communication (3343)
   (a) Cytokine (2548)
   (b) Chemotactic (111)

#### S4 Describing trained models for functional biological activity classification of peptide sequendes

98 functional biological activity classification models were trained using a combination of supervised learning algorithms and pre-trained models. Table S3 shows statistical descriptors of the performances obtained for the trained models.

Table S3: Statistical descriptions of the performances obtained for the trained models

| | **Accuracy** | **F1-score** | **Precision** | **Recall** | **MCC** |
|---|---|---|---|---|---|
| **mean** | 0.873 | 0.873 | 0.877 | 0.873 | 0.750 |
| **std** | 0.054 | 0.055 | 0.053 | 0.054 | 0.108 |
| **min** | 0.753 | 0.753 | 0.754 | 0.753 | 0.505 |
| **25%** | 0.835 | 0.835 | 0.844 | 0.835 | 0.677 |
| **50%** | 0.875 | 0.875 | 0.879 | 0.875 | 0.756 |
| **75%** | 0.903 | 0.903 | 0.910 | 0.903 | 0.815 |
| **max** | 1.0 | 1.0 | 1.0 | 1.0 | 1.0 |

Table S4 summarizes all trained models' sensitivity and specificity performances.

Table S4: Statistical descriptors for sensitivity and specificity for the trained classification models

| **Descriptor** | **Sensitivity** | **Specificity** |
|---|---|---|
| Mean | 0.877 | 0.873 |
| STD | 0.053 | 0.054 |
| Min | 0.754 | 0.753 |
| 25% | 0.844 | 0.835 |
| 50% | 0.879 | 0.875 |
| 75% | 0.910 | 0.903 |
| Max | 1.000 | 1.000 |

Figure S1 summarises the distribution performances by encoders obtained during the validation process for all trained functional biological activity classification models.

Figure S2 summarises the distribution performances by algorithms obtained during the validation process for all trained functional biological activity classification models.

Figure S3 summarises the frequency of the encoders and algorithms selected for trained binary classification models. In the case of encoders, Prottrans t5 XLU50, Prottrans UniRef, and Prottrans T5BDF are selected in more than 70% of the trained models. In contrast, algorithms like ExtraTrees and Gaussian Process are selected in more than 50% of the trained models.

Table S5 summarises the frequency of combinations of algorithms and encoders selected for trained functional biological classification models.

Table S5: Number of selected combinations of algorithm and encoder by selected, trained model for functional biological activity classification

| **Algorithm** | **Encoder** | **Activity** |
|---|---|---|
| ExtraTrees | ProTrans UniRef | 9 |
| GaussianProcess | ProTrans UniRef | 8 |
| ExtraTrees | ProTrans XLU50 | 7 |
| ExtraTrees | ESM1b | 6 |
| KNeighbors | ProTrans XLU50 | 5 |
| KNeighbors | ProTrans UniRef | 5 |
| ExtraTrees | ProTrans T5BDF | 5 |

| | | |
|---|---|---|
| GaussianProcess | ProTrans XLU50 | 4 |
| GaussianProcess | ESM1b | 4 |
| GaussianProcess | ProTrans T5BDF | 4 |
| RandomForest | ProTrans T5BDF | 4 |
| XGB | ProTrans XLU50 | 4 |
| GradientBoosting | ESM1b | 3 |
| GradientBoosting | ProTrans UniRef | 2 |
| KNeighbors | ProTrans T5BDF | 2 |
| RandomForest | ProTrans UniRef | 2 |
| RandomForest | ProTrans XLU50 | 2 |
| XGB | ProTrans T5BDF | 2 |
| GradientBoosting | ProTrans T5BDF | 2 |
| XGB | ProTrans XLNET | 2 |
| ExtraTrees | ProTrans ALBERT | 2 |
| Bagging | prottrans_bert | 2 |
| AdaBoost | ProTrans XLNET | 1 |
| ExtraTrees | ProTrans XLNET | 1 |
| ExtraTrees | prottrans_bert | 1 |
| RandomForest | ESM1b | 1 |
| RandomForest | ProTrans ALBERT | 1 |
| Bagging | ProTrans T5BDF | 1 |
| Bagging | ProTrans XLU50 | 1 |
| Bagging | ProTrans UniRef | 1 |
| XGB | ESM1b | 1 |
| XGB | ProTrans UniRef | 1 |
| Bagging | ESM1b | 1 |
| AdaBoost | ESM1b | 1 |

Table S6 summarizes the 98 trained classification models by reporting the sensitivity and the specificity performances.

Table S6: Summary of sensitivity and specificity for the 98 binary classification models trained in Peptipedia v2.0

| **Activity** | **Algorithm** | **Pre-trained model** | **Sensitivity** | **Specificity** |
|---|---|---|---|---|
| Activator | ExtraTrees | ProTrans t5 xlu50 | 0.85 | 0.84 |
| Allergen | GaussianProcess | ProTrans t5 Uniref | 0.91 | 0.91 |
| Angiotensin-converting enzyme (ace) inhibitors | GaussianProcess | esm1b | 0.89 | 0.88 |
| Angiotensinase inhibitor | KNeighbors | ProTrans t5 xlu50 | 0.89 | 0.89 |
| Anti SARS-COV | ExtraTrees | ProTrans t5 Uniref | 0.91 | 0.91 |
| Anti SARS-COV 2 | GradientBoosting | ProTrans t5-BDF | 0.87 | 0.86 |
| Anti aging | ExtraTrees | ProTrans t5 Uniref | 0.86 | 0.86 |
| Anti amnesic | ExtraTrees | ProTrans t5-ALBERT | 0.91 | 0.9 |
| Anti andes virus | RandomForest | ProTrans t5-BDF | 0.98 | 0.98 |
| *Anti bunyaviridae* | XGB | ProTrans t5-BDF | 0.99 | 0.99 |
| *Anti coronaviridae* | RandomForest | ProTrans t5-ALBERT | 0.85 | 0.84 |
| Anti diabetic | GaussianProcess | ProTrans t5-BDF | 0.86 | 0.85 |
| Anti diabetic type 1 | ExtraTrees | ProTrans t5 Uniref | 0.89 | 0.89 |
| Anti diabetic type 2 | GradientBoosting | ProTrans t5 Uniref | 0.89 | 0.89 |
| Anti feline immunodeficiency virus | ExtraTrees | esm1b | 0.91 | 0.9 |

| | | | | |
|---|---|---|---|---|
| *Anti filoviridae* | ExtraTrees | esm1b | 0.86 | 0.86 |
| *Anti flaviviridae* | ExtraTrees | ProTrans t5 xlu50 | 0.88 | 0.87 |
| *Anti hantaviridae* | AdaBoost | ProTrans t5-XLNET | 0.98 | 0.98 |
| Anti hepatitis B virus | Bagging | ProTrans t5-BERT | 0.88 | 0.88 |
| Anti hepatitis C virus | ExtraTrees | ProTrans t5-ALBERT | 0.9 | 0.89 |
| Anti herpes simplex virus | ExtraTrees | ProTrans t5 Uniref | 0.87 | 0.87 |
| Anti herpesviridae | ExtraTrees | ProTrans t5 Uniref | 0.81 | 0.81 |
| Anti human parainfluenza virus | RandomForest | ProTrans t5-BDF | 1.0 | 1.0 |
| Anti hypertensive | KNeighbors | ProTrans t5 Uniref | 0.9 | 0.9 |
| Anti inflamatory | GaussianProcess | ProTrans t5 Uniref | 0.88 | 0.88 |
| Anti nematode | ExtraTrees | ProTrans t5 xlu50 | 0.94 | 0.93 |
| *Anti orthomyxoviridae* | GradientBoosting | esm1b | 0.84 | 0.84 |
| Anti oxidative | XGB | ProTrans t5-BDF | 0.85 | 0.85 |
| *Anti paramyxoviridae* | XGB | ProTrans t5-XLNET | 0.91 | 0.91 |
| Anti poxviridae | ExtraTrees | esm1b | 0.93 | 0.92 |
| Anti respiratory syncytial virus | XGB | ProTrans t5 xlu50 | 0.94 | 0.94 |
| Anti sin nombre virus | RandomForest | ProTrans t5-BDF | 0.99 | 0.99 |
| Anti thrombotic | RandomForest | ProTrans t5 Uniref | 0.89 | 0.88 |
| Anti vaccinia | Bagging | ProTrans t5-BERT | 0.95 | 0.94 |
| Anuran defense | ExtraTrees | esm1b | 0.89 | 0.89 |
| Cell cell communication | ExtraTrees | ProTrans t5 xlu50 | 0.89 | 0.89 |
| Cell penetrating | GaussianProcess | ProTrans t5-BDF | 0.85 | 0.85 |
| Coagulation + vascular | KNeighbors | ProTrans t5 Uniref | 0.87 | 0.87 |
| Cytokine | GaussianProcess | ProTrans t5 Uniref | 0.96 | 0.96 |
| DNA-binding | Bagging | ProTrans t5 Uniref | 0.99 | 0.99 |
| Drug delivery vehicle | GaussianProcess | ProTrans t5 Uniref | 0.81 | 0.81 |
| Enzyme activator | ExtraTrees | ProTrans t5-BDF | 0.85 | 0.85 |
| Enzyme inhibitor | GaussianProcess | ProTrans t5 xlu50 | 0.88 | 0.88 |
| Glucose | GaussianProcess | ProTrans t5-BDF | 0.86 | 0.85 |
| Gluten immunogenic + celiac toxic | XGB | ProTrans t5-XLNET | 0.99 | 0.99 |
| Hormone | GaussianProcess | ProTrans t5 xlu50 | 0.88 | 0.88 |
| Immunological | GaussianProcess | ProTrans t5 xlu50 | 0.86 | 0.86 |
| Immunomodulatory | KNeighbors | ProTrans t5-BDF | 0.85 | 0.84 |
| Immunoregulator | Bagging | ProTrans t5-BDF | 0.82 | 0.81 |
| Inhibitor | GaussianProcess | esm1b | 0.88 | 0.88 |
| Insecticidal | GaussianProcess | esm1b | 0.86 | 0.86 |
| Metabolic | GaussianProcess | ProTrans t5 Uniref | 0.81 | 0.8 |
| Molecular binding | ExtraTrees | esm1b | 0.9 | 0.87 |
| Neurological | XGB | esm1b | 0.84 | 0.84 |
| Neuropeptide | GaussianProcess | ProTrans t5 Uniref | 0.86 | 0.85 |
| Neurotoxin | GradientBoosting | esm1b | 0.97 | 0.97 |
| Opioid | ExtraTrees | ProTrans t5-BDF | 0.95 | 0.94 |
| Pain related | RandomForest | esm1b | 0.93 | 0.93 |
| Potentiator | ExtraTrees | ProTrans t5-BDF | 0.92 | 0.92 |
| Propeptide | XGB | ProTrans t5 xlu50 | 0.88 | 0.88 |
| Quorum sensing | ExtraTrees | ProTrans t5-BERT | 0.84 | 0.84 |
| Regulatory | KNeighbors | ProTrans t5 xlu50 | 0.83 | 0.83 |
| Signal peptide | KNeighbors | ProTrans t5 xlu50 | 0.88 | 0.88 |
| Taste | GradientBoosting | ProTrans t5-BDF | 0.95 | 0.95 |
| Transit | KNeighbors | ProTrans t5 xlu50 | 0.97 | 0.97 |

| | | | | |
|---|---|---|---|---|
| Tumor homing | RandomForest | ProTrans t5 xlu50 | 0.92 | 0.91 |
| Tumor targeting | RandomForest | ProTrans t5 xlu50 | 0.89 | 0.89 |
| Venous | XGB | ProTrans t5 xlu50 | 0.92 | 0.9 |
| Anti angiogenic | ExtraTrees | ProTrans t5 Uniref | 0.79 | 0.79 |
| Anti biofilm | ExtraTrees | esm1b | 0.79 | 0.79 |
| Anti candida | ExtraTrees | ProTrans t5 xlu50 | 0.75 | 0.75 |
| Anti coronavirus | ExtraTrees | ProTrans t5 Uniref | 0.81 | 0.8 |
| Anti dengue virus | Bagging | ProTrans t5 xlu50 | 0.88 | 0.83 |
| Anti endotoxin | ExtraTrees | ProTrans t5 xlu50 | 0.82 | 0.81 |
| Anti HIV | GaussianProcess | ProTrans t5 Uniref | 0.82 | 0.82 |
| Anti influenza virus | ExtraTrees | ProTrans t5-XLNET | 0.79 | 0.79 |
| *Anti retroviridae* | GaussianProcess | ProTrans t5 xlu50 | 0.8 | 0.8 |
| Antiprotozoal | GaussianProcess | ProTrans t5-BDF | 0.82 | 0.81 |
| Blood brain barrier penetrating | GradientBoosting | ProTrans t5 Uniref | 0.8 | 0.8 |
| Chemotactic | ExtraTrees | ProTrans t5 xlu50 | 0.88 | 0.88 |
| Cosmetic and dermatology | RandomForest | ProTrans t5-BDF | 0.79 | 0.78 |
| Cytotoxic | GaussianProcess | ProTrans t5 Uniref | 0.8 | 0.8 |
| Hemolytic | GaussianProcess | esm1b | 0.84 | 0.83 |
| Pheromone | ExtraTrees | ProTrans t5-BDF | 0.89 | 0.89 |
| Protease inhibitor | AdaBoost | esm1b | 0.86 | 0.86 |
| Stimulator | GradientBoosting | esm1b | 0.85 | 0.85 |
| Anti gram (-) | KNeighbors | ProTrans t5-BDF | 0.81 | 0.81 |
| Anti gram (+) | XGB | ProTrans t5 xlu50 | 0.89 | 0.89 |
| Antibacterial | ExtraTrees | ProTrans t5 Uniref | 0.89 | 0.89 |
| Anticancer | KNeighbors | ProTrans t5 Uniref | 0.81 | 0.8 |
| Anti fungal | ExtraTrees | ProTrans t5 Uniref | 0.92 | 0.92 |
| Antimicrobial | KNeighbors | ProTrans t5 Uniref | 0.88 | 0.88 |
| Anti tuberculosis | ExtraTrees | ProTrans t5-BDF | 0.85 | 0.83 |
| Antiviral | KNeighbors | ProTrans t5 xlu50 | 0.86 | 0.86 |
| Therapeutic | KNeighbors | ProTrans t5 Uniref | 0.88 | 0.88 |
| Toxic | XGB | ProTrans t5 Uniref | 0.93 | 0.93 |
| Antiparasitic | RandomForest | ProTrans t5 Uniref | 0.81 | 0.81 |
| *Anti methicillin-resistant S. aureus* | Bagging | esm1b | 0.81 | 0.81 |

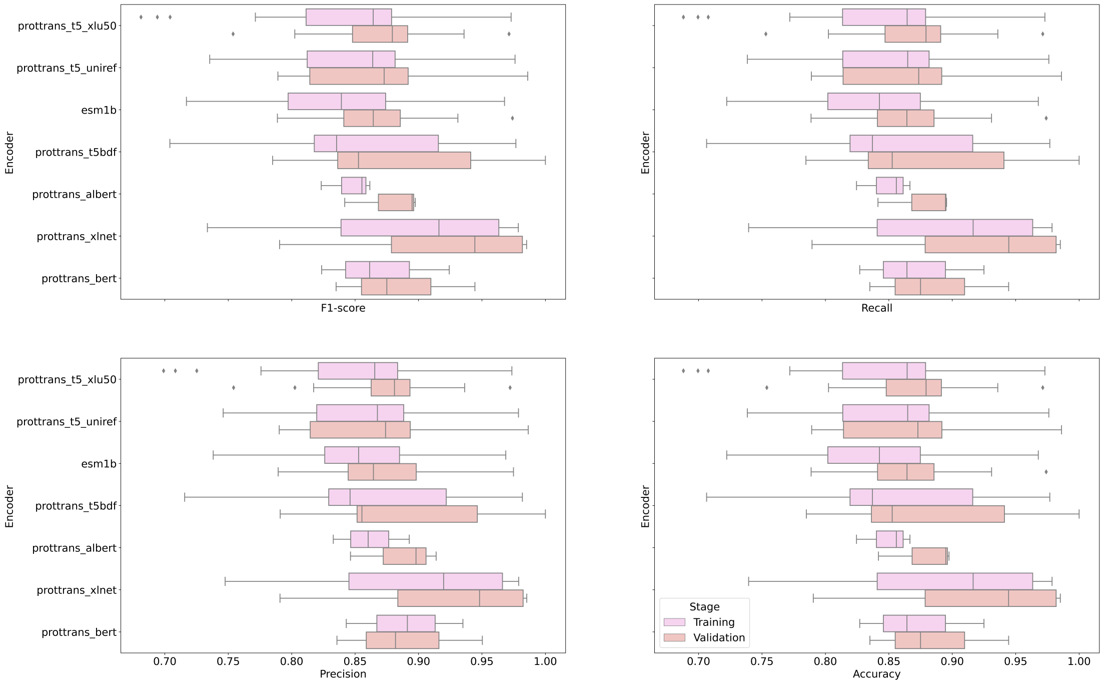


Supplementary Figure S1: Statistical distribution performances by encoders for the trained models, including the precision, recall, F1-score, and accuracy.

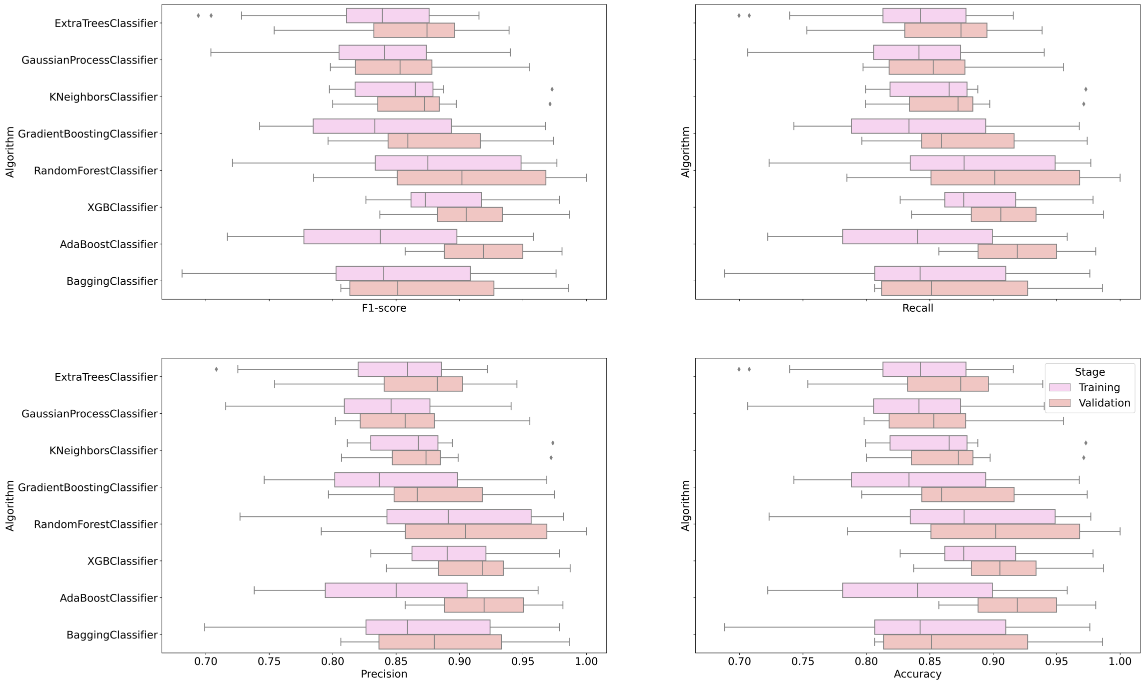


Supplementary Figure S2: Statistical distribution performances by algorithms for the trained models, including the precision, recall, F1-score, and accuracy.

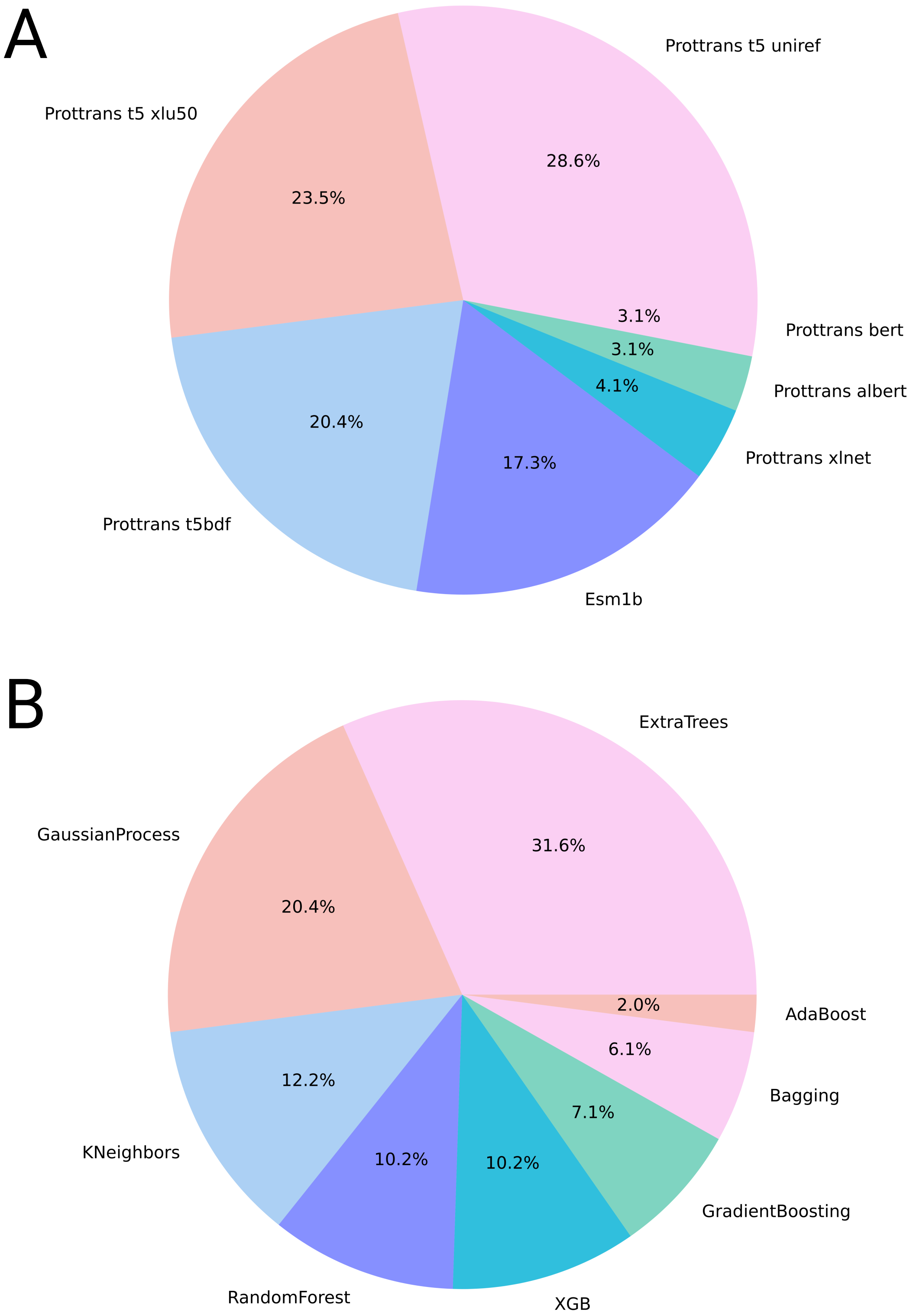


Supplementary Figure S3: Summary of the frequency of encoders (A) and algorithms (B) selected for generating the trained binary classification models.

### S5 Summary of the new main functionalities implemented on Peptipedia v2.0

Peptipedia v2.0 has implemented a new face for its user-friendly web platform. Figure S4 summarises the homepage available on Peptipedia v2.0.

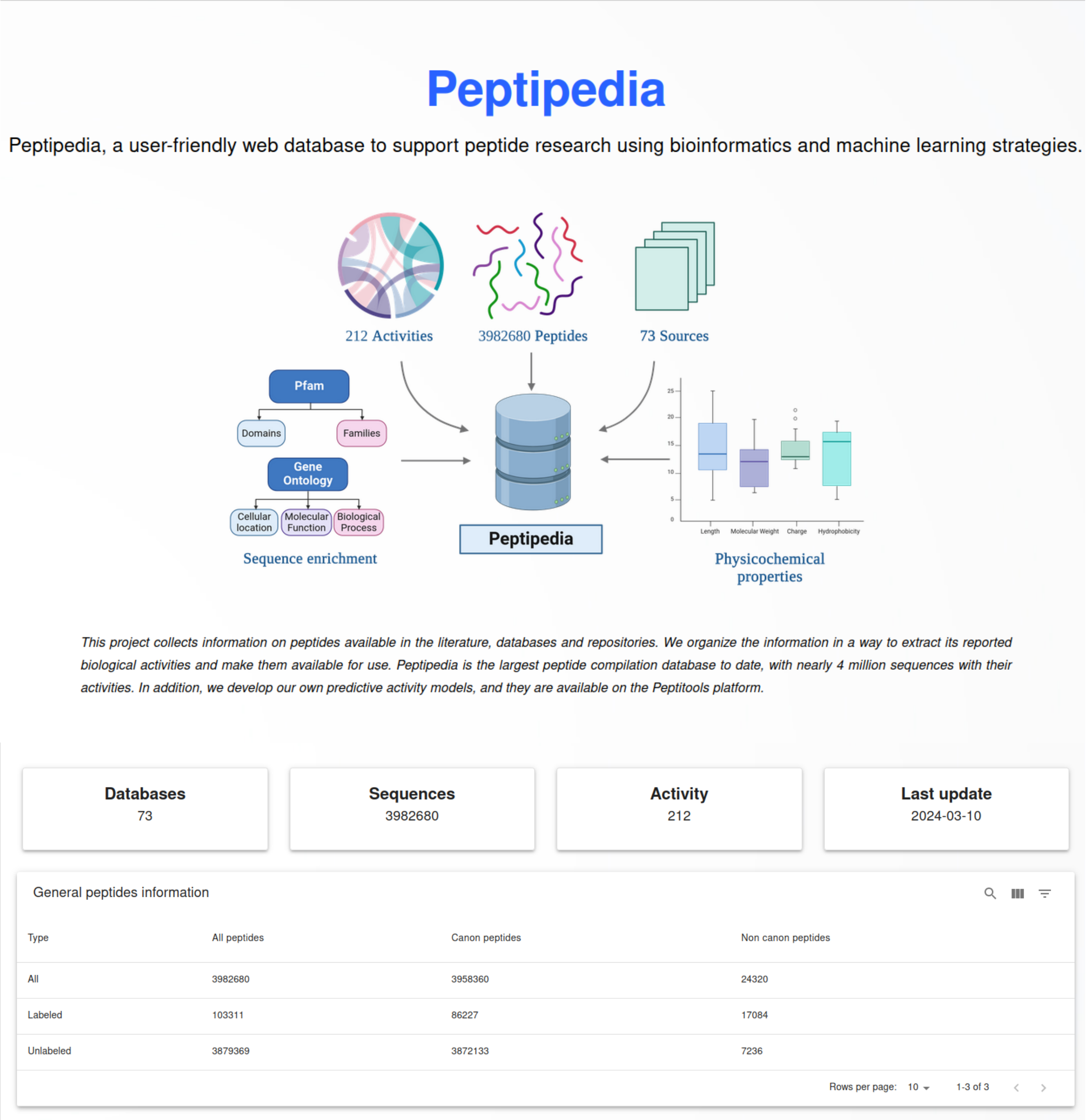


Supplementary Figure S4: Home page implemented for Peptipedia v2.0, including a schematic representation of the implemented workflow to process data and the statistics for the collected peptides.

The activities in Peptipedia v2.0 are listed in a tree format, representing the functional biological activity tree explained in the previous section and updated in this new version of Peptipedia v2.0. Figure S5 summarises the activity visualization.

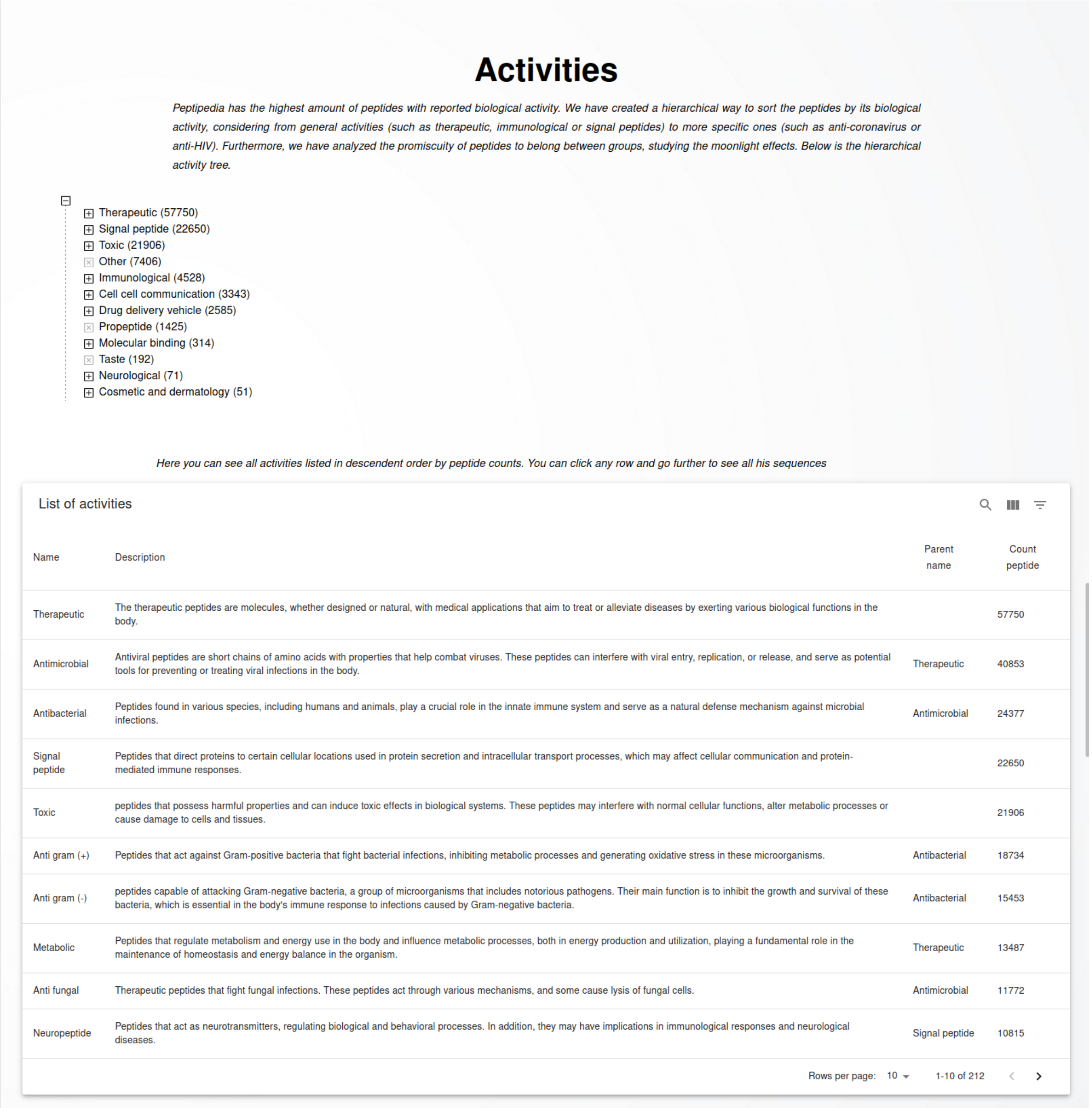

### Activities

*Peptipedia has the highest amount of peptides with reported biological activity. We have created a hierarchical way to sort the peptides by its biological activity, considering from general activities (such as therapeutic, immunological or signal peptides) to more specific ones (such as anti-coronavirus or anti-HIV). Furthermore, we have analyzed the promiscuity of peptides to belong between groups, studying the moonlight effects. Below is the hierarchical activity tree.*

- Therapeutic (57750)
- Signal peptide (22650)
- Toxic (21906)
- Other (7406)
- Immunological (4528)
- Cell cell communication (3343)
- Drug delivery vehicle (2585)
- Propeptide (1425)
- Molecular binding (314)
- Taste (192)
- Neurological (71)
- Cosmetic and dermatology (51)

*Here you can see all activities listed in descendent order by peptide counts. You can click any row and go further to see all his sequences*

List of activities

| Name | Description | Parent name | Count peptide |
|---|---|---|---|
| Therapeutic | The therapeutic peptides are molecules, whether designed or natural, with medical applications that aim to treat or alleviate diseases by exerting various biological functions in the body. | | 57750 |
| Antimicrobial | Antiviral peptides are short chains of amino acids with properties that help combat viruses. These peptides can interfere with viral entry, replication, or release, and serve as potential tools for preventing or treating viral infections in the body. | Therapeutic | 40853 |
| Antibacterial | Peptides found in various species, including humans and animals, play a crucial role in the innate immune system and serve as a natural defense mechanism against microbial infections. | Antimicrobial | 24377 |
| Signal peptide | Peptides that direct proteins to certain cellular locations used in protein secretion and intracellular transport processes, which may affect cellular communication and protein-mediated immune responses. | | 22650 |
| Toxic | peptides that possess harmful properties and can induce toxic effects in biological systems. These peptides may interfere with normal cellular functions, alter metabolic processes or cause damage to cells and tissues. | | 21906 |
| Anti gram (+) | Peptides that act against Gram-positive bacteria that fight bacterial infections, inhibiting metabolic processes and generating oxidative stress in these microorganisms. | Antibacterial | 18734 |
| Anti gram (-) | peptides capable of attacking Gram-negative bacteria, a group of microorganisms that includes notorious pathogens. Their main function is to inhibit the growth and survival of these bacteria, which is essential in the body's immune response to infections caused by Gram-negative bacteria. | Antibacterial | 15453 |
| Metabolic | Peptides that regulate metabolism and energy use in the body and influence metabolic processes, both in energy production and utilization, playing a fundamental role in the maintenance of homeostasis and energy balance in the organism. | Therapeutic | 13487 |
| Anti fungal | Therapeutic peptides that fight fungal infections. These peptides act through various mechanisms, and some cause lysis of fungal cells. | Antimicrobial | 11772 |
| Neuropeptide | Peptides that act as neurotransmitters, regulating biological and behavioral processes. In addition, they may have implications in immunological responses and neurological diseases. | Signal peptide | 10815 |

Rows per page: 10 1-10 of 212

Supplementary Figure S5: Visualization of functional biological activities collected and reported in Peptipedia v2.0. On top, a tree visualization is deployed. On the bottom is a description of each activity, including the name, full description, and number of sequences for each activity, differentiating by the reported peptide in literature and predictive activity

The new version of the sequence search engine in Peptipedia v2.0 facilitates the application of different filters to customize the type of information identification. Query results have also been optimized through the generation of materialized views. The visualization of the results has been updated to present the characteristics of the identified sequences in a more user-friendly manner, incorporating all existing information in the platform (see Figure S6).

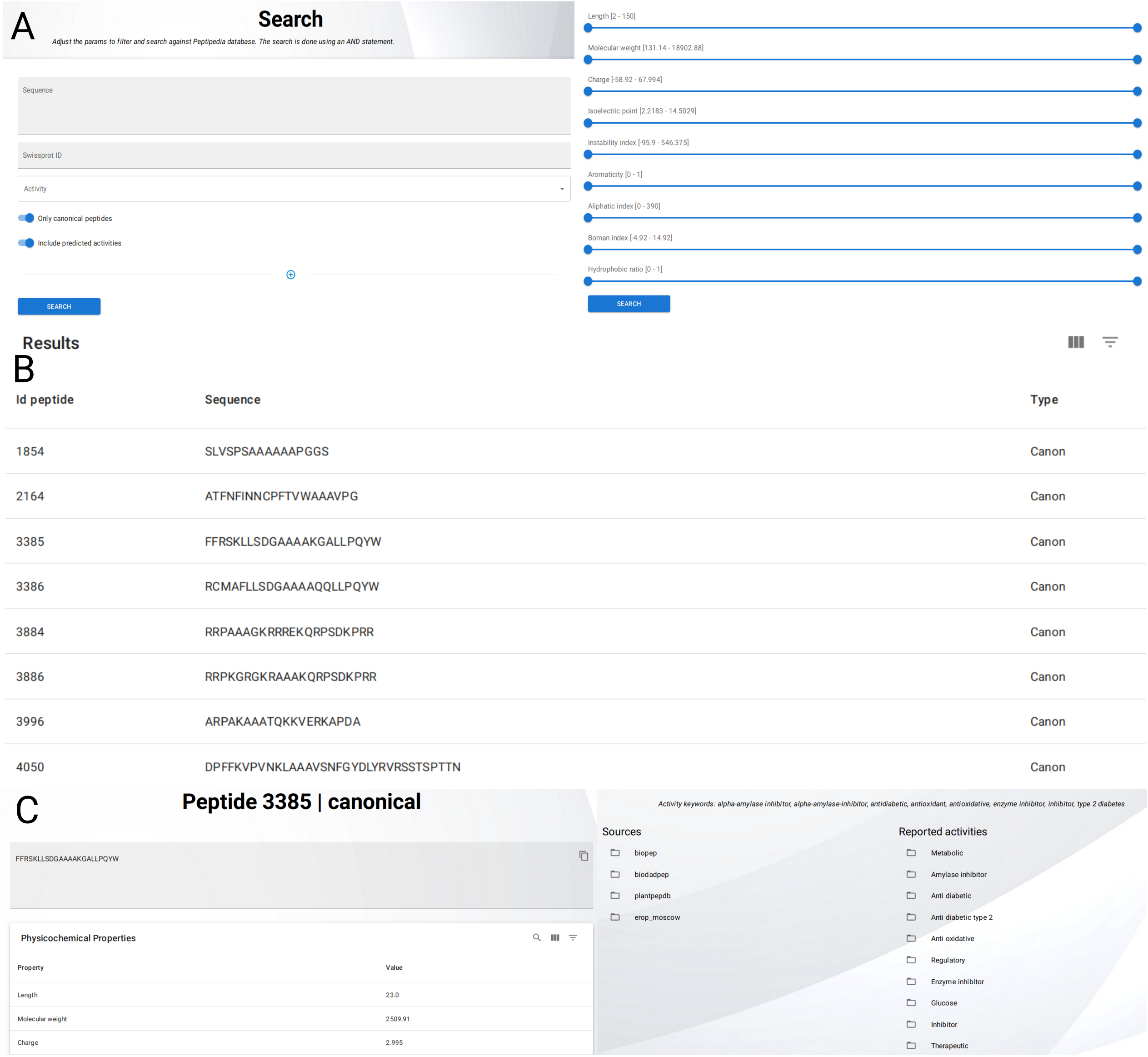


Supplementary Figure S6: **Schematic representation of the search menu and results available on Peptipedia v2.0**. **A** Schematic representation of search tool. Peptipedia v2.0 makes the search by pattern available, and it is also possible to apply different filters, including physicochemical properties and select desirable activities. **B** Once the search is processed, the results are displayed as a table showing the peptide ID, peptide sequences and the peptide category based on the canonical analysis. **C** The user can access a full detail for each identified peptide by clicking on any row in the Table. The full details include the sequence, status of the paper, different biological activities, data sources, related keywords, physicochemical and thermodynamic characterization, function domain, Gene Ontology, and secondary structure prediction.

The supervised learning system in Peptipetia facilitates the design and training of predictive models using classic supervised learning algorithms. Figure S7 summarises the training configuration, metric performances, and evaluation process. First, the configuration is required to train a model. These configurations include i) selecting the task and configuration of the test dataset size and $k-$fold validation process, ii) algorithm selection, iii) numerical representation strategy, and iv) applying standardization or dimensional reduction techniques. Once the configuration is processed, the model is trained and delivered. The tool displays different metrics to evaluate the predictive model's performance, including classic metrics, a learning curve, and a sensitivity-specificity analysis in the classification task. In contrast, scatter plots are displayed to evaluate the prediction error for the regression task. Finally, the trained model can be used to assess new peptide sequences. The sequences need to be uploaded, the same configurations for preprocessing sequences will be applied, and the trained model will be used to generate the desirable predictions. The predictions are displayed in the tool and are available to download.

The functionality of the clustering tool is similar to that of the supervised learning tool. Figure S8 summarises the clustering process, including the configuration process, the metrics to evaluate the performance of the algorithms, and the groups' visualization.

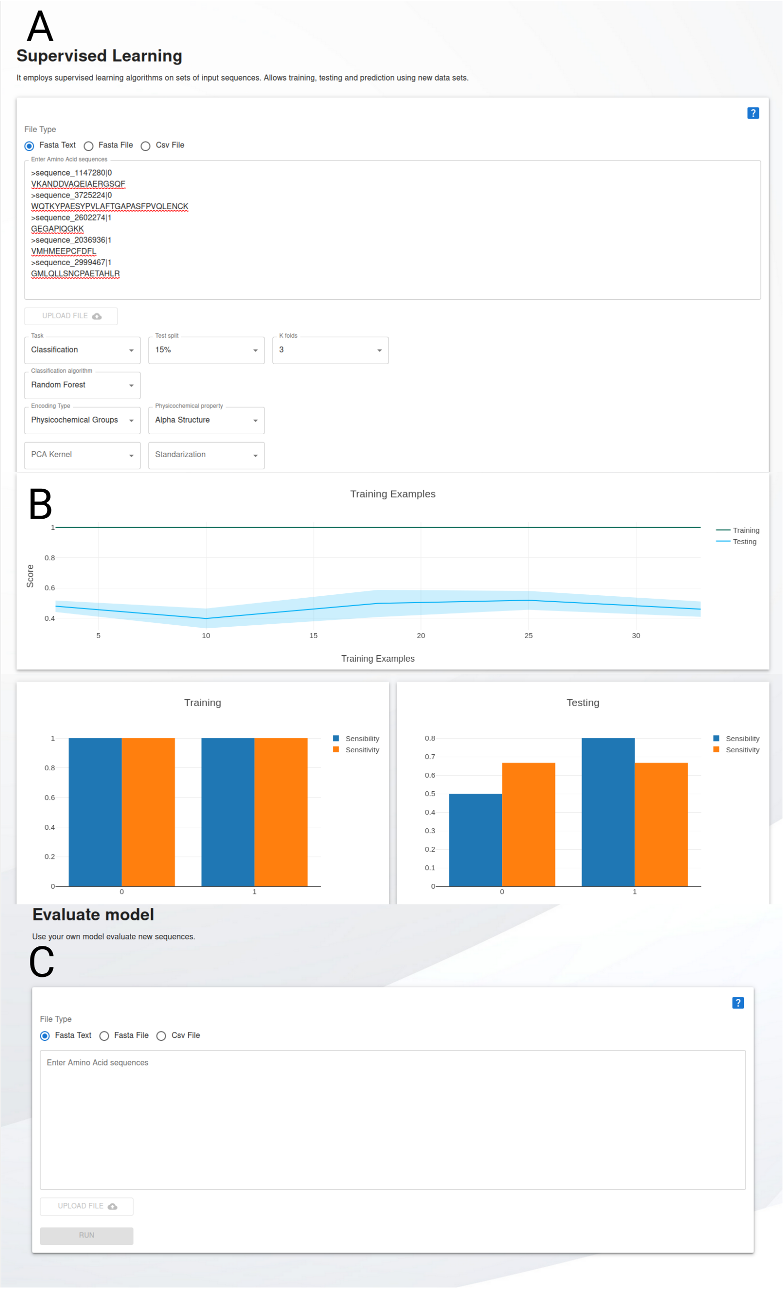


Supplementary Figure S7: **A schematic representation of the training process using the supervised learning service available in Peptipedia v2.0**. **A** Configurations for the training process. The configurations include i) selecting the task and configuration of the test dataset size and $k$−fold validation process, ii) algorithm selection, iii) numerical representation strategy, and iv) applying standardization or dimensional reduction techniques. **B** Summary of the main results generated post-training process. The results include i) a summary table with all performances, ii) a learning curve plot and a confusion matrix, and iii) an analysis of sensitivity and specificity values. **C** The trained model can be used with new peptide sequences employing the evaluation system. In this tool, the model is used to predict peptide sequences added by the user. Once the sequences are loaded, the tool processes the sequences, applying the same configurations employed to train the model, and then the model makes the predictions. The predictions are displayed in the tool.

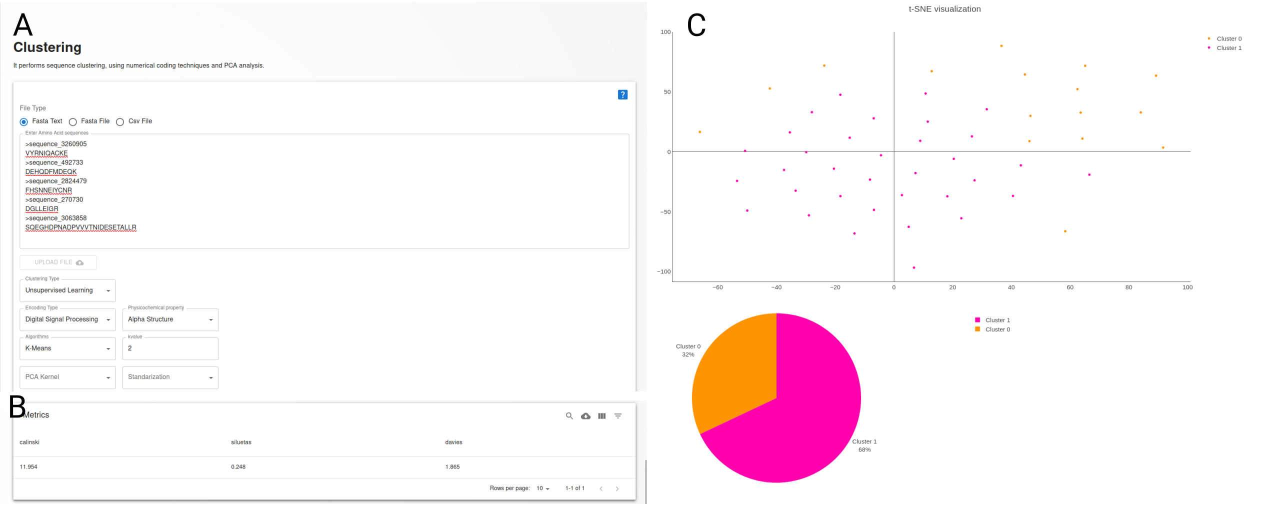


Supplementary Figure S8: **Clustering service available in Peptipedia v2.0**. **A** The configuration tool necessary to apply the clustering process. **B** The result generated by the clustering process focused on the metrics. **C** The groups' visualizations by t-SNE analysis and piechart.

### References

[1] Piyush Agrawal, Dhruv Bhagat, Manish Mahalwal, Neelam Sharma, and Gajendra P S Raghava. AntiCP 2.0: an updated model for predicting anticancer peptides. *Briefings in Bioinformatics*, 22(3), August 2020.

[2] Piyush Agrawal, Sherry Bhalla, Kumardeep Chaudhary, Rajesh Kumar, Meenu Sharma, and Gajendra P. S. Raghava. In silico approach for prediction of antifungal peptides. *Frontiers in Microbiology*, 9, February 2018.

[3] Sajid Ahmed, Rafsanjani Muhammod, Zahid Hossain Khan, Sheikh Adilina, Alok Sharma, Swakkhar Shatabda, and Abdollah Dehzangi. ACP-MHCNN: an accurate multi-headed deep-convolutional neural network to predict anticancer peptides. *Scientific Reports*, 11(1), December 2021.

[4] Plaimein Amnuaycheewa, Mohamed Abdelmoteleb, John Wise, Barbara Bohle, Fatima Ferreira, Afua O. Tetteh, Steve L. Taylor, and Richard E. Goodman. Development of a sequence searchable database of celiac disease-associated peptides and proteins for risk assessment of novel food proteins. *Frontiers in Allergy*, 3, May 2022.

[5] Data Analysis & Modeling Group at Hasselt University, Functional Genomics, and Proteomics Unit at K.U. Leuven. Peptidedb: Bioactive peptide database. `http://www.peptides.be/`, 30 Apr, 2008. Accessed: 2023-10-16.

[6] Amos Bairoch and Rolf Apweiler. The swiss-prot protein sequence database and its supplement trembl in 2000. *Nucleic acids research*, 28(1):45–48, 2000.

[7] Yannan Bin, Wei Zhang, Wending Tang, Ruyu Dai, Menglu Li, Qizhi Zhu, and Junfeng Xia. Prediction of neuropeptides from sequence information using ensemble classifier and hybrid features. *Journal of Proteome Research*, 19(9):3732–3740, August 2020.

[8] Michał Burdukiewicz, Katarzyna Sidorczuk, Dominik Rafacz, Filip Pietluch, Mateusz Bakała, Jadwiga Słowik, and Przemysław Gagat. CancerGram: An effective classifier for differentiating anticancer from antimicrobial peptides. *Pharmaceutics*, 12(11):1045, October 2020.

[9] Stephen K Burley, Helen M Berman, Cole Christie, Jose M Duarte, Zukang Feng, John Westbrook, Jasmine Young, and Christine Zardecki. Rcsb protein data bank: Sustaining a living digital data resource

that enables breakthroughs in scientific research and biomedical education. *Protein Science*, 27(1):316–330, 2018.

[10] Anita Chaudhary, Sherry Bhalla, Sumeet Patiyal, Gajendra P.S. Raghava, and Girish Sahni. Fermfoodb: A database of bioactive peptides derived from fermented foods. *Heliyon*, 7(4):e06668, April 2021.

[11] Kumardeep Chaudhary, Ritesh Kumar, Sandeep Singh, Abhishek Tuknait, Ankur Gautam, Deepika Mathur, Priya Anand, Grish C. Varshney, and Gajendra P. S. Raghava. A web server and mobile app for computing hemolytic potency of peptides. *Scientific Reports*, 6(1), March 2016.

[12] Chia-Ru Chung, Ting-Rung Kuo, Li-Ching Wu, Tzong-Yi Lee, and Jorng-Tzong Horng. Characterization and identification of antimicrobial peptides with different functional activities. *Briefings in Bioinformatics*, 21(3):1098–1114, 06 2019.

[13] Ruyu Dai, Wei Zhang, Wending Tang, Evelien Wynendaele, Qizhi Zhu, Yannan Bin, Bart De Spiegeleer, and Junfeng Xia. BBPpred: Sequence-based prediction of blood-brain barrier peptides with feature representation learning and logistic regression. *Journal of Chemical Information and Modeling*, 61(1):525–534, January 2021.

[14] Christian Dallago, Konstantin Schütze, Michael Heinzinger, Tobias Olenyi, Maria Littmann, Amy X. Lu, Kevin K. Yang, Seonwoo Min, Sungroh Yoon, James T. Morton, and Burkhard Rost. Learned embeddings from deep learning to visualize and predict protein sets. *Current Protocols*, 1(5):e113, 2021.

[15] Durdam Das, Mohini Jaiswal, Fatima Nazish Khan, Shahzaib Ahamad, and Shailesh Kumar. PlantPepDB: A manually curated plant peptide database. *Scientific Reports*, 10(1), February 2020.

[16] Frank Desiere, Eric W Deutsch, Nichole L King, Alexey I Nesvizhskii, Parag Mallick, Jimmy Eng, Sharon Chen, James Eddes, Sandra N Loevenich, and Ruedi Aebersold. The peptideatlas project. *Nucleic acids research*, 34(suppl_1):D655–D658, 2006.

[17] Sylvia Van Dorpe, Antoon Bronselaer, Joachim Nielandt, Sofie Stalmans, Evelien Wynendaele, Kurt Audenaert, Christophe Van De Wiele, Christian Burvenich, Kathelijne Peremans, Hung Hsuchou, Guy De Tré, and Bart De Spiegeleer. Brainpeps: the blood–brain barrier peptide database. *Brain Structure and Function*, 217(3):687–718, December 2011.

[18] Kassel & thpr.net e. K. Dr. Katja Kapp. Signal peptide website: An information platform for signal sequences and signal peptides. `http://www.signalpeptide.de/`, 11 Jun, 2010. Accessed: 2023-10-16.

[19] Ahmed Elnaggar, Michael Heinzinger, Christian Dallago, Ghalia Rehawi, Yu Wang, Llion Jones, Tom Gibbs, Tamas Feher, Christoph Angerer, Martin Steinegger, et al. Prottrans: Toward understanding the language of life through self-supervised learning. *IEEE transactions on pattern analysis and machine intelligence*, 44(10):7112–7127, 2021.

[20] Xiaodan Fan, Lu Bai, Liang Zhu, Li Yang, and Xuewu Zhang. Marine algae-derived bioactive peptides for human nutrition and health. *Journal of Agricultural and Food Chemistry*, 62(38):9211–9222, September 2014.

[21] Ankur Gautam, Kumardeep Chaudhary, Rahul Kumar, Arun Sharma, Pallavi Kapoor, Atul Tyagi, and Gajendra P S Raghava and. In silico approaches for designing highly effective cell penetrating peptides. *Journal of Translational Medicine*, 11(1), March 2013.

[22] Ankur Gautam, Kumardeep Chaudhary, Sandeep Singh, Anshika Joshi, Priya Anand, Abhishek Tuknait, Deepika Mathur, Grish C. Varshney, and Gajendra P. S. Raghava. Hemolytik: a database of experimentally determined hemolytic and non-hemolytic peptides. *Nucleic Acids Research*, 42(D1):D444–D449, October 2013.

[23] Richard E. Goodman, Motohiro Ebisawa, Fatima Ferreira, Hugh A. Sampson, Ronald van Ree, Stefan Vieths, Joseph L. Baumert, Barbara Bohle, Sreedevi Lalithambika, John Wise, and Steve L. Taylor. AllergenOnline: A peer-reviewed, curated allergen database to assess novel food proteins for potential cross-reactivity. *Molecular Nutrition &amp Food Research*, 60(5):1183–1198, March 2016.

[24] Sudheer Gupta, Pallavi Kapoor, Kumardeep Chaudhary, Ankur Gautam, Rahul Kumar, and Gajendra P. S. Raghava and. In silico approach for predicting toxicity of peptides and proteins. *PLoS ONE*, 8(9):e73957, September 2013.

[25] Riadh Hammami, Abdelmajid Zouhir, Jeannette Ben Hamida, and Ismail Fliss. BACTIBASE: a new web-accessible database for bacteriocin characterization. *BMC Microbiology*, 7(1), October 2007.

[26] Jhih-Hua Jhong, Lantian Yao, Yuxuan Pang, Zhongyan Li, Chia-Ru Chung, Rulan Wang, Shangfu Li, Wenshuo Li, Mengqi Luo, Renfei Ma, Yuqi Huang, Xiaoning Zhu, Jiahong Zhang, Hexiang Feng, Qifan Cheng, Chunxuan Wang, Kun Xi, Li-Ching Wu, Tzu-Hao Chang, Jorng-Tzong Horng, Lizhe Zhu, Ying-Chih Chiang, Zhuo Wang, and Tzong-Yi Lee. dbAMP 2.0: updated resource for antimicrobial peptides with an enhanced scanning method for genomic and proteomic data. *Nucleic Acids Research*, 50(D1):D460–D470, November 2021.

[27] Q. Kaas, R. Yu, A.-H. Jin, S. Dutertre, and D. J. Craik. ConoServer: updated content, knowledge, and discovery tools in the conopeptide database. *Nucleic Acids Research*, 40(D1):D325–D330, November 2011.

[28] Xinyue Kang, Fanyi Dong, Cheng Shi, Shicai Liu, Jian Sun, Jiaxin Chen, Haiqi Li, Hanmei Xu, Xingzhen Lao, and Heng Zheng. DRAMP 2.0, an updated data repository of antimicrobial peptides. *Scientific Data*, 6(1), August 2019.

[29] Pallavi Kapoor, Harinder Singh, Ankur Gautam, Kumardeep Chaudhary, Rahul Kumar, and Gajendra P. S. Raghava. TumorHoPe: A database of tumor homing peptides. *PLoS ONE*, 7(4):e35187, April 2012.

[30] Kimia Kardani and Azam Bolhassani. Cppsite 2.0: An available database of experimentally validated cell-penetrating peptides predicting their secondary and tertiary structures. *Journal of Molecular Biology*, 433(11):166703, May 2021.

[31] Mst. Shamima Khatun, Md. Mehedi Hasan, and Hiroyuki Kurata. PreAIP: Computational prediction of anti-inflammatory peptides by integrating multiple complementary features. *Frontiers in Genetics*, 10, March 2019.

[32] Yoona Kim, Steven Bark, Vivian Hook, and Nuno Bandeira. NeuroPedia: neuropeptide database and spectral library. *Bioinformatics*, 27(19):2772–2773, August 2011.

[33] Ravi Kumar, Kumardeep Chaudhary, Minakshi Sharma, Gandharva Nagpal, Jagat Singh Chauhan, Sandeep Singh, Ankur Gautam, and Gajendra P.S. Raghava. AHTPDB: a comprehensive platform for analysis and presentation of antihypertensive peptides. *Nucleic Acids Research*, 43(D1):D956–D962, November 2014.

[34] Vinod Kumar, Sumeet Patiyal, Anjali Dhall, Neelam Sharma, and Gajendra Pal Singh Raghava. B3pred: A random-forest-based method for predicting and designing blood–brain barrier penetrating peptides. *Pharmaceutics*, 13(8):1237, August 2021.

[35] Travis J Lawrence, Dana L Carper, Margaret K Spangler, Alyssa A Carrell, Tomás A Rush, Stephen J Minter, David J Weston, and Jessy L Labbé. amPEPpy 1.0: a portable and accurate antimicrobial peptide prediction tool. *Bioinformatics*, 37(14):2058–2060, 11 2020.

[36] Hao-Ting Lee, Chen-Che Lee, Je-Ruei Yang, Jim Z. C. Lai, and Kuan Y. Chang. A large-scale structural classification of antimicrobial peptides. *BioMed Research International*, 2015:1–6, 2015.

[37] Supatcha Lertampaiporn, Tayvich Vorapreeda, Apiradee Hongsthong, and Chinae Thammarongtham. Ensemble-AMPPred: Robust AMP prediction and recognition using the ensemble learning method with a new hybrid feature for differentiating AMPs. *Genes*, 12(2):137, January 2021.

[38] Chenkai Li, Darcy Sutherland, S. Austin Hammond, Chen Yang, Figali Taho, Lauren Bergman, Simon Houston, René L. Warren, Titus Wong, Linda M. N. Hoang, Caroline E. Cameron, Caren C. Helbing, and Inanc Birol. AMPlify: attentive deep learning model for discovery of novel antimicrobial peptides effective against WHO priority pathogens. *BMC Genomics*, 23(1), January 2022.

[39] Yanchao Liu, Youzhuo Zhu, Xin Sun, Tianyue Ma, Xingzhen Lao, and Heng Zheng. Dravp: A comprehensive database of antiviral peptides and proteins. *Viruses*, 15(4):820, March 2023.

[40] Mariagrazia Di Luca, Giuseppe Maccari, Giuseppantonio Maisetta, and Giovanna Batoni. BaAMPs: the database of biofilm-active antimicrobial peptides. *Biofouling*, 31(2):193–199, February 2015.

[41] Zhibin Lv, Feifei Cui, Quan Zou, Lichao Zhang, and Lei Xu. Anticancer peptides prediction with deep representation learning features. *Briefings in Bioinformatics*, 22(5), February 2021.

[42] Deepika Mathur, Satya Prakash, Priya Anand, Harpreet Kaur, Piyush Agrawal, Ayesha Mehta, Rajesh Kumar, Sandeep Singh, and Gajendra P. S. Raghava. Peplife: A repository of the half-life of peptides. *Scientific Reports*, 6(1), November 2016.

[43] Deepika Mathur, Sandeep Singh, Ayesha Mehta, Piyush Agrawal, and Gajendra P. S. Raghava. In silico approaches for predicting the half-life of natural and modified peptides in blood. *PLOS ONE*, 13(6):e0196829, June 2018.

[44] Minkiewicz, Iwaniak, and Darewicz. BIOPEP-UWM database of bioactive peptides: Current opportunities. *International Journal of Molecular Sciences*, 20(23):5978, November 2019.

[45] Gandharva Nagpal, Kumardeep Chaudhary, Piyush Agrawal, and Gajendra P. S. Raghava. Computer-aided prediction of antigen presenting cell modulators for designing peptide-based vaccine adjuvants. *Journal of Translational Medicine*, 16(1), July 2018.

[46] Mario Novković, Juraj Simunić, Viktor Bojović, Alessandro Tossi, and Davor Juretić. DADP: the database of anuran defense peptides. *Bioinformatics*, 28(10):1406–1407, March 2012.

[47] Poonam Pandey, Vinal Patel, Nithin V. George, and Sairam S. Mallajosyula. KELM-CPPpred: Kernel extreme learning machine based prediction model for cell-penetrating peptides. *Journal of Proteome Research*, 17(9):3214–3222, July 2018.

[48] Yuxuan Pang, Lantian Yao, Jhih-Hua Jhong, Zhuo Wang, and Tzong-Yi Lee. Avpiden: a new scheme for identification and functional prediction of antiviral peptides based on machine learning approaches. *Briefings in Bioinformatics*, 22(6), July 2021.

[49] Sandy S Pineda, Pierre-Alain Chaumeil, Anne Kunert, Quentin Kaas, Mike W C Thang, Lien Le, Michael Nuhn, Volker Herzig, Natalie J Saez, Ben Cristofori-Armstrong, Raveendra Anangi, Sebastian Senff, Dominique Gorse, and Glenn F King. ArachnoServer 3.0: an online resource for automated discovery, analysis and annotation of spider toxins. *Bioinformatics*, 34(6):1074–1076, October 2017.

[50] Stefano P. Piotto, Lucia Sessa, Simona Concilio, and Pio Iannelli. YADAMP: yet another database of antimicrobial peptides. *International Journal of Antimicrobial Agents*, 39(4):346–351, April 2012.

[51] Malak Pirtskhalava, Anthony A Amstrong, Maia Grigolava, Mindia Chubinidze, Evgenia Alimbarashvili, Boris Vishnepolsky, Andrei Gabrielian, Alex Rosenthal, Darrell E Hurt, and Michael Tartakovsky. DBAASP v3: database of antimicrobial/cytotoxic activity and structure of peptides as a resource for development of new therapeutics. *Nucleic Acids Research*, 49(D1):D288–D297, November 2020.

[52] Abid Qureshi, Himani Tandon, and Manoj Kumar. AVP-ICsub50/subpred: Multiple machine learning techniques-based prediction of peptide antiviral activity in terms of half maximal inhibitory concentration (ICsub50/sub). *Peptide Science*, 104(6):753–763, November 2015.

[53] Abid Qureshi, Nishant Thakur, and Manoj Kumar. HIPdb: A database of experimentally validated HIV inhibiting peptides. *PLoS ONE*, 8(1):e54908, January 2013.

[54] Abid Qureshi, Nishant Thakur, Himani Tandon, and Manoj Kumar. AVPdb: a database of experimentally validated antiviral peptides targeting medically important viruses. *Nucleic Acids Research*, 42(D1):D1147–D1153, November 2013.

[55] Kunjulakshmi R, Ambuj Kumar, Keerthana Vinod Kumar, Avik Sengupta, Kavita Kundal, Simran Sharma, Ankita Pawar, Pithani Sai Krishna, Mohammad Alfatah, Sandipan Ray, Bhavana Tiwari, and Rahul Kumar. Aagingbase: a comprehensive database of anti-aging peptides. *Database*, 2024, January 2024.

[56] Akanksha Rajput, Amit Kumar Gupta, and Manoj Kumar. Prediction and analysis of quorum sensing peptides based on sequence features. *PLOS ONE*, 10(3):e0120066, March 2015.

[57] Azhagiya Singam Ettayapuram Ramaprasad, Sandeep Singh, Raghava Gajendra P. S, and Subramanian Venkatesan. AntiAngioPred: A server for prediction of anti-angiogenic peptides. *PLOS ONE*, 10(9):e0136990, September 2015.

[58] Github Repository. Identification of antimicrobial peptides using spaced-seeds and reduced amino-acid alphabets. `https://github.com/gmit-amp-res/amp`, 27 Jun, 2021. Accessed: 2023-10-16.

[59] Susanta Roy. BioDADPep: A bioinformatics database for anti-diabetic peptides. *Bioinformation*, 15(11):780–783, November 2019.

[60] Sandeep Singh, Kumardeep Chaudhary, Sandeep Kumar Dhanda, Sherry Bhalla, Salman Sadullah Usmani, Ankur Gautam, Abhishek Tuknait, Piyush Agrawal, Deepika Mathur, and Gajendra P.S. Raghava. SATPdb: a database of structurally annotated therapeutic peptides. *Nucleic Acids Research*, 44(D1):D1119–D1126, November 2015.

[61] Xin Su, Jing Xu, Yanbin Yin, Xiongwen Quan, and Han Zhang. Antimicrobial peptide identification using multi-scale convolutional network. *BMC Bioinformatics*, 20(1), December 2019.

[62] Zhelyazko Terziyski, Margarita Terziyska, Ivelina Deseva, Stanka Hadzhikoleva, Albert Krastanov, Dasha Mihaylova, and Emil Hadzhikolev. Peplab platform: Database and software tools for analysis of food-derived bioactive peptides. *Applied Sciences*, 13(2):961, January 2023.

[63] Patrick Brendan Timmons and Chandralal M Hewage. ENNAVIA is a novel method which employs neural networks for antiviral and anti-coronavirus activity prediction for therapeutic peptides. *Briefings in Bioinformatics*, 22(6), July 2021.

[64] Atul Tyagi, Pallavi Kapoor, Rahul Kumar, Kumardeep Chaudhary, Ankur Gautam, and G. P. S. Raghava. In silico models for designing and discovering novel anticancer peptides. *Scientific Reports*, 3(1), October 2013.

[65] Atul Tyagi, Abhishek Tuknait, Priya Anand, Sudheer Gupta, Minakshi Sharma, Deepika Mathur, Anshika Joshi, Sandeep Singh, Ankur Gautam, and Gajendra P.S. Raghava. CancerPPD: a database of anticancer peptides and proteins. *Nucleic Acids Research*, 43(D1):D837–D843, September 2014.

[66] Salman Sadullah Usmani, Rajesh Kumar, Vinod Kumar, Sandeep Singh, and Gajendra P S Raghava. Antitbpdb: a knowledgebase of anti-tubercular peptides. *Database*, 2018, January 2018.

[67] Ronald van Ree, Dexter Sapiter Ballerda, M. Cecilia Berin, Laurent Beuf, Alexander Chang, Gabriele Gadermaier, Paul A. Guevera, Karin Hoffmann-Sommergruber, Emir Islamovic, Liisa Koski, John Kough, Gregory S. Ladics, Scott McClain, Kyle A. McKillop, Shermaine Mitchell-Ryan, Clare A. Narrod, Lucilia Pereira Mouriès, Syril Pettit, Lars K. Poulsen, Andre Silvanovich, Ping Song, Suzanne S. Teuber, and Christal Bowman. The COMPARE database: A public resource for allergen identification, adapted for continuous improvement. *Frontiers in Allergy*, 2, August 2021.

[68] Daniel Veltri, Uday Kamath, and Amarda Shehu. Deep learning improves antimicrobial peptide recognition. *Bioinformatics*, 34(16):2740–2747, March 2018.

[69] Faiza Hanif Waghu, Ram Shankar Barai, Pratima Gurung, and Susan Idicula-Thomas. CAMPsubr3/sub: a database on sequences, structures and signatures of antimicrobial peptides: Table 1. *Nucleic Acids Research*, 44(D1):D1094–D1097, October 2015.

[70] Guangshun Wang, Xia Li, and Zhe Wang. APD3: the antimicrobial peptide database as a tool for research and education. *Nucleic Acids Research*, 44(D1):D1087–D1093, November 2015.

[71] Jian Wang, Tailang Yin, Xuwen Xiao, Dan He, Zhidong Xue, Xinnong Jiang, and Yan Wang. StraPep: a structure database of bioactive peptides. *Database*, 2018, January 2018.

[72] Evelien Wynendaele, Antoon Bronselaer, Joachim Nielandt, Matthias D'Hondt, Sofie Stalmans, Nathalie Bracke, Frederick Verbeke, Christophe Van De Wiele, Guy De Tré, and Bart De Spiegeleer. Quorumpeps database: chemical space, microbial origin and functionality of quorum sensing peptides. *Nucleic Acids Research*, 41(D1):D655–D659, November 2012.

[73] Guizi Ye, Hongyu Wu, Jinjiang Huang, Wei Wang, Kuikui Ge, Guodong Li, Jiang Zhong, and Qingshan Huang. LAMP2: a major update of the database linking antimicrobial peptides. *Database*, 2020, January 2020.

[74] Hai-Cheng Yi, Zhu-Hong You, Xi Zhou, Li Cheng, Xiao Li, Tong-Hai Jiang, and Zhan-Heng Chen. ACP-DL: A deep learning long short-term memory model to predict anticancer peptides using high-efficiency feature representation. *Molecular Therapy - Nucleic Acids*, 17:1–9, September 2019.

[75] Lezheng Yu, Runyu Jing, Fengjuan Liu, Jiesi Luo, and Yizhou Li. DeepACP: A novel computational approach for accurate identification of anticancer peptides by deep learning algorithm. *Molecular Therapy - Nucleic Acids*, 22:862–870, December 2020.

[76] A. A. Zamyatnin. The EROP-moscow oligopeptide database. *Nucleic Acids Research*, 34(90001):D261–D266, January 2006.

[77] Jinhao Zhang, Zehua Zhang, Lianrong Pu, Jijun Tang, and Fei Guo. AIEpred: An ensemble predictive model of classifier chain to identify anti-inflammatory peptides. *IEEE/ACM Transactions on Computational Biology and Bioinformatics*, 18(5):1831–1840, September 2021.

[78] Qianyue Zhang, Xue Chen, Bowen Li, Chunying Lu, Shanshan Yang, Jinjin Long, Heng Chen, Jian Huang, and Bifang He. A database of anti-coronavirus peptides. *Scientific Data*, 9(1), June 2022.
